# Gene networks are conserved across reproductive development between the fern *Ceratopteris richardii* and the flowering plant *Arabidopsis thaliana*

**DOI:** 10.1101/2025.03.18.643782

**Authors:** Andrew R.G. Plackett, Marco Catoni

## Abstract

How plants first invented seeds is a longstanding unresolved evolutionary question. Seed-bearing plants arose from within seedless vascular plants, with ferns as their closest relatives, but how the seed developmental programme first originated has remained intractable through comparative morphology or the fossil record. To investigate this question at the level of gene network evolution, we established a transcriptional expression atlas across sporophyte and gametophyte reproductive development of the fern *Ceratopteris richardii* and compared reproductive-associated genes to those of *Arabidopsis thaliana*. Conservation was detected between distinct organs. As in flowering, the hormone gibberellin promoted sporing of the fern sporophyte shoot. Sporophyll genes were conserved with floral development. Gametophyte post-fertilization egg chamber (archegonium) genes were conserved with both pre- and post-fertilization seed development, as were two post-fertilization signals, auxin and sugar. We conclude that the seed may first have arisen through the mis-expression of archegonium developmental programmes within the developing sporophyte sporangium.

## INTRODUCTION

The evolution of the seed entailed dramatic changes to plant reproduction, arising from a single origin within the seedless vascular plants^1,2^ and with monilophytes (horsetails and ferns) the closest surviving outgroup^2,3,4^. Extant and extinct seeds (‘ovules’ before fertilization) share a common body-plan: a diploid nucellus apically specialised for pollen reception, containing a multicellular haploid egg-producing structure, with a subtending chalaza from which one or more integuments arise^5^. Ovule tissues undergo further development after fertilization to form the seed coat and (in angiosperms) specialised storage tissues (the endosperm)^6,7,8^. But how this developmentally-complex organ first originated is a longstanding unresolved question.

Seed evolution has been deduced by comparing to ancestral seedless reproduction still found in extant ferns, in which separate free-living diploid and haploid ‘generations’ both form reproductive organs^9,10^. The diploid sporophyte forms reproductively-modified fronds (sporophylls) within which meiosis generates haploid spores. The thalloid gametophytes that germinate from these produce gametangia through mitosis: antheridia (containing free-swimming sperm) and archegonia (egg chambers) which when fertilized undergo post-fertilization modifications to support embryo development^11^. Early ovule evolution is proposed to have involved sporophyll modifications to encapsulate a reduced gametophyte (endospory)^12,13^, with the archegonium still present in gymnosperms but lost in angiosperms^7^. A further crucial innovation was heterospory, genetically-predetermined male and female sporangia (pollen sac and ovule nucellus) arising from ancestral homosporous sporangia^14,15,16,17^. Many ferns remain homosporous, including the genetic model *Ceratopteris richardii*^18^. ‘Male’ (micro)sporogenesis resembles ancestral development whereas ‘female’ (mega)sporogenesis has apparently undergone significant modification^19^, now typically resulting in a single viable megaspore (monospory)^20,21^. The genetic changes underlying these evolutionary transitions remain unknown.

A complete molecular hypothesis for ovule evolution has not been proposed, but a role for the MADS-box gene family, which specifies angiosperm ovule identity^22,23^ has long been suggested^24,25^. Recent analysis of individual fern genes supports the partial conservation of reproductive gene networks^24,26,27,28^, but to what extent is unclear. Although fern genomic information is increasingly available^29,30,31,32,33^, as are transcriptomes at either whole-lifecycle resolution^34,35,36^ or focussed on single reproductive organs^30,37^, a detailed characterisation of gene networks across fern reproductive development to compare to seed plant reproduction is lacking. We thus undertook mRNA-seq analysis to identify genes associated with reproductive processes in both the *Ceratopteris* sporophyte and gametophyte. We identified and experimentally validated a conserved role for the phytohormone gibberellin (GA) in the sporophyte shoot reproductive transition and two post-fertilization regulatory mechanisms between the seed and the fern archegonium. GO term enrichment and whole-network comparison suggest that ovule/seed gene networks are conserved specifically with the post-fertilization fern archegonium. This raises a new hypothesis that the ovule arose by mis-expressing post-fertilization archegonium gene networks within the sporophyte.

## RESULTS

### Identifying reproductive gene networks in the *Ceratopteris* sporophyte

To establish a tractable experimental system, the *Ceratopteris* sporophyte reproductive transition was first characterised. The shoot apex switched to reproductive development (‘sporing’) consistently between replicates, chronologically (100% by 18 weeks) and developmentally (20.6 ± 0.07 fronds) (**Extended Data Fig. 1; Supplementary Data 1**). Transcriptional changes associated with reproductive development were identified by comparing gene expression across frond development between the sporing phase (130 days old) (**Fig. 1a-f**) and the vegetative phase immediately prior to sporing (60 days old) (**Fig. 1g-l**). Further 25-day-old apex and primordia samples were included as a control for the sporing transition (**Supplementary** Fig. 1). Frond development was divided into sequential stages common to vegetative fronds and sporophylls: the shoot apical region (‘Apx’) (**Fig. 1b,h**); frond primordium (‘Prm’) (**Fig. 1c,i; Supplementary** Fig. 1); fiddlehead (‘Fid’) (**Fig. 1d,j**); expanding frond (‘Exp’) (**Fig. 1e,k**) and mature frond (‘Mat’) (**Fig. 1f,l**). Sporangia were visible by the start of sporophyll expansion (**Extended Data Fig. 2a-l**), with sporangium sizes suggesting overlapping development throughout (**Extended Data Fig. 2m,n**). Post-meiotic sporangia were only observed in mature sporophyll samples (**Extended Data Fig. 2e-h**).

**Fig. 1.**
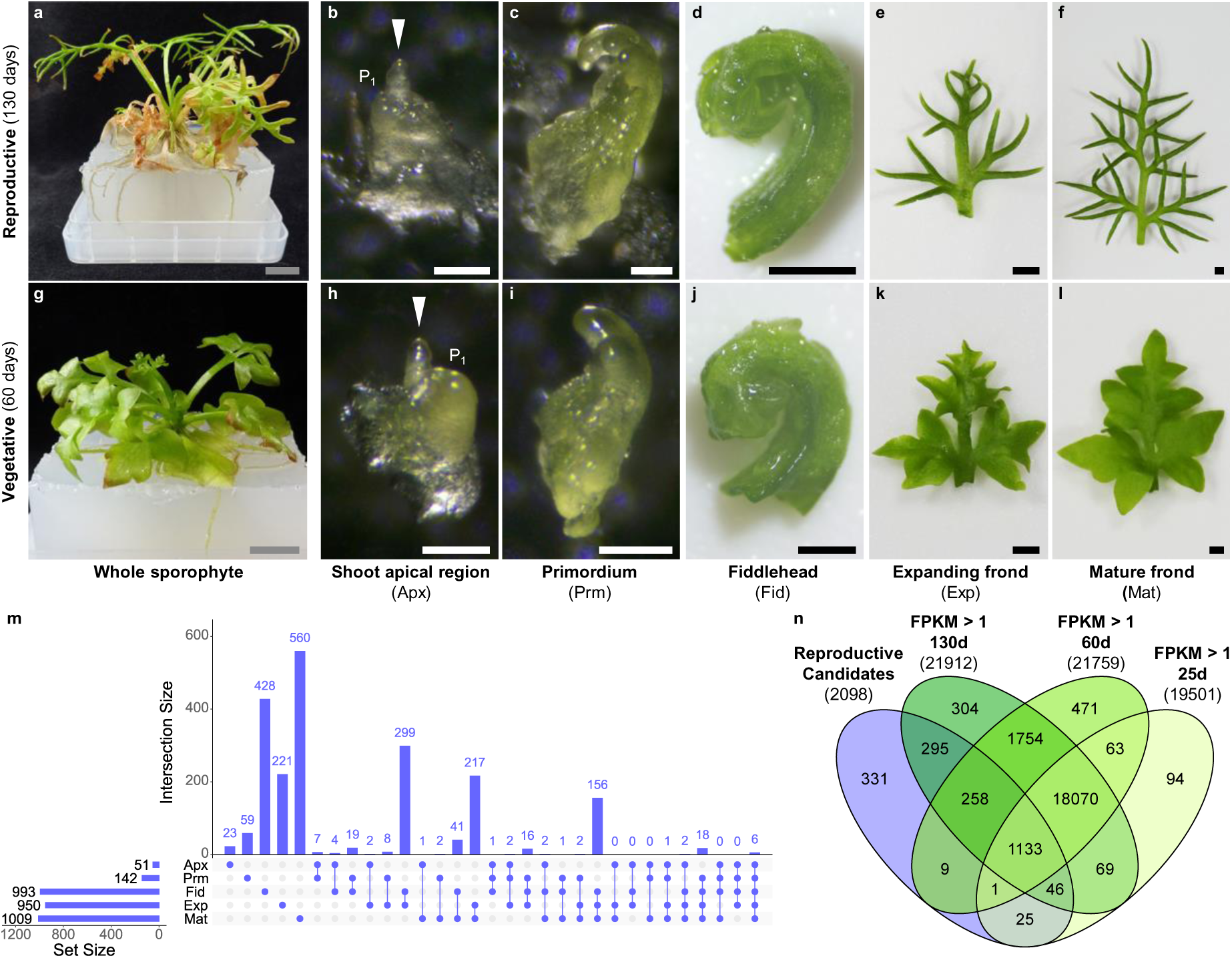
Reproduction-associated genes were identified across *Ceratopteris* sporophyll development. **a-l**, The sporing *Ceratopteris* shoot (**a**) was dissected into sequential samples across frond development: the shoot apical region (Apx) (**b**) comprising the shoot apex (white arrowhead) and early primordia (P1), the next-most recent primordium (Prm) (**c**) undergoing morphogenesis, the frond fiddlehead (Fid) (**d**) emerged from the shoot apical region, expanding fronds (Exp) (**e**) where the fiddlehead had terminated, from unfurling to approximately 75% of maturity, and mature non-senescent fronds (Mat) (**f**) where growth had ceased compared to prior fronds. Candidate reproductive genes were identified through significantly increased abundance (Padj < 0.05, log2fold expression change > 1) in sporophyll samples vs corresponding frond samples in the vegetative phase immediately preceding the sporing transition (**g-l**). Scale bars = 10 mm (grey), 200 µm (white) and 1 mm (black). **m**,**n** Comparing reproductive candidate reproductive genes between frond developmental stages identified 2098 candidates (**m**). These were categorised based on transcript abundance (**n**): candidates at FPKM >1 at 130d but not in vegetative phases as ‘sporophyll-specific’; at FPKM >1 in both 130d and at least one vegetative phase as ‘sporophyll-enriched’; and at FPKM <1 at 130d as ‘sporophyll-FPKM<1’.

Principal Component Analysis (PCA) and mRNA-seq expression correlation confirmed that biological replicates clustered by sample (**Extended Data Fig. 3a**, **Supplementary** Fig. 2). Frond developmental stages separated progressively along PC1 (excluding Apx and Prm, which overlapped except for 25d Prm) while vegetative and reproductive phases separated across PC2. Differential expression (DE) analysis identified 2098 transcripts with increased abundance (Padj < 0.05, log2foldchange >1) in sporophylls vs vegetative controls in at least one frond stage (**Fig. 1m; Supplementary Data 2**). Among these we categorised 295 as present (FPKM >1) only in the sporophyll (‘sporophyll-specific’) and 1437 present in the sporophyll and at least one vegetative stage (‘sporophyll-enriched’) (**Fig. 1n**), representing potential direct regulators of reproduction and genes either with dual vegetative and reproductive function or frond developmental genes whose expression is modified in sporophylls. 366 remaining candidates were expressed at FPKM <1 in all sporophyll samples.

### Gibberellin promotes the Ceratopteris sporing transition

Because Apx and Prm samples overlapped under PCA, Apx reproductive candidates were analysed separately (**Fig. 2a**). Fifteen significantly-enriched reproductive GO terms (p < 0.05) were identified and all associated with one MADS-box gene, *CMADS1*^24^ (**Supplementary Data 3**). Genes promoting a shoot reproductive transition were predicted to have increased expression immediately prior to it, and to be preferentially expressed in the shoot apex vs primordia. Expression data was used to rank thirteen Apx candidates with progressive significant increases in abundance (Padj < 0.05) between 25d, 60d and 130d (**Fig. 2b**). These included *CMADS1*, but this was present in all ages and was not preferential to Apx samples (Padj > 0.05). Apx candidates contained two further MADS-box homologs which showed increased abundance at 130d only (**Supplementary Data 2**).

**Fig. 2.**
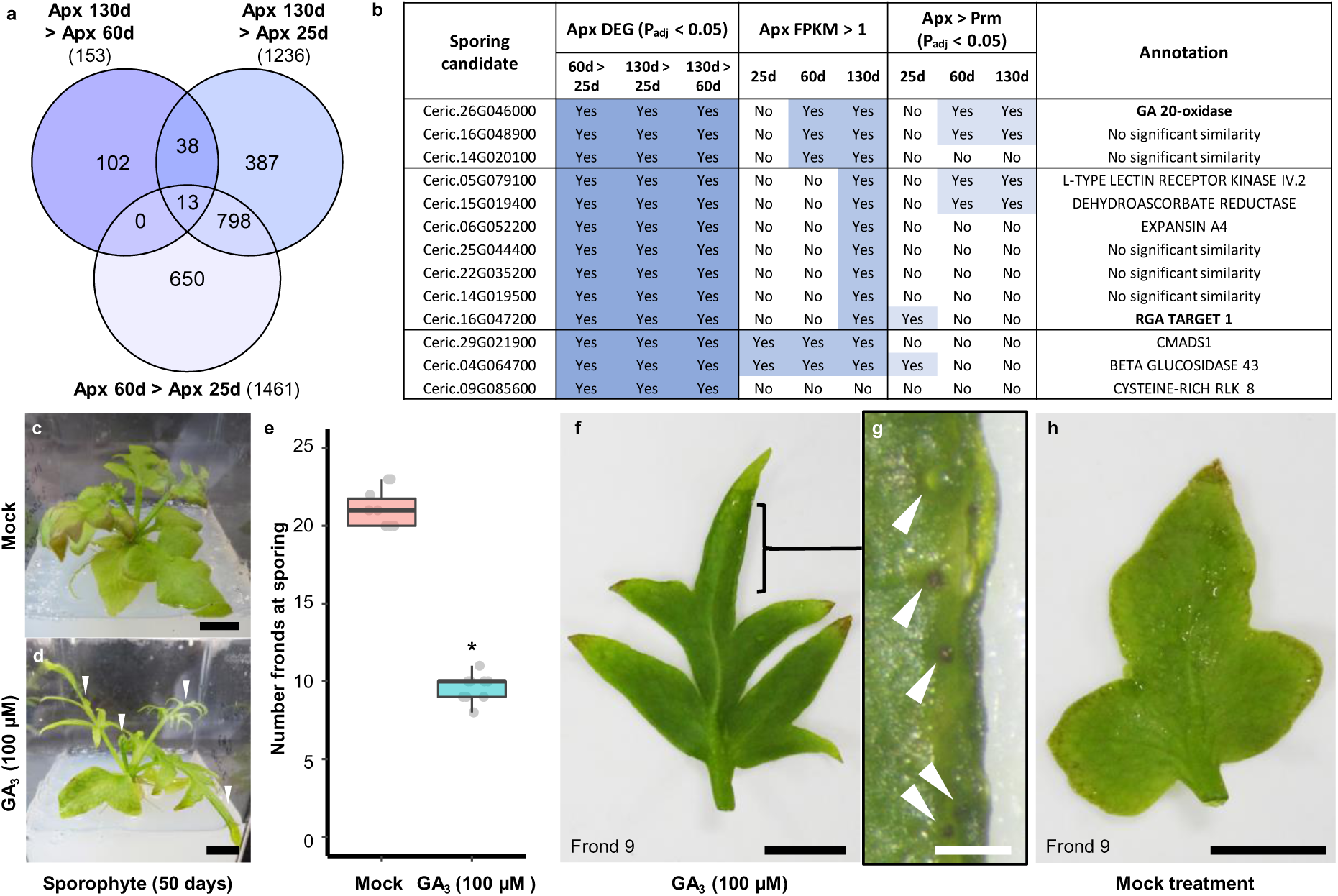
Gibberellin promotes the *Ceratopteris* sporophyte shoot sporing transition. **a**, Differential gene expression analysis identified 51 candidates with significantly greater expression in the 130d Apx vs 60d and 25d, with 13 showing a progressive increase immediately prior to the sporing transition (**a**). These 13 candidates were ranked according to their similarity to the predicted expression characteristics of a shoot apex sporing regulator (**b**), with the strongest candidate being a GA 20-oxidase gibberellin biosynthesis gene. **c-h**, Under exogenous GA treatment the sporing transition of the *Ceratopteris* sporophyte is accelerated both chronologically (**c,d**) and developmentally (**e**). The first spore-bearing fronds under GA treatment (**f**) exhibit margin rolling and sporangium production (**g**) compared to mock-treated fronds at the same position (**h**). White arrowheads denote visible sporophylls (**d**) and sporangia within the rolled frond margin (**g**). Asterisk denotes a significant difference (p < 0.05) between mock and GA treatment. *n* = 11. Scale bars = 10 mm (black), 500 µm (white). Pairwise comparisons were performed using two-tailed Mann-Whitney tests (**Supplementary Data 1**).

Candidates whose expression profile matched that predicted for sporing regulators included a GA 20-oxidase GA biosynthetic enzyme (**Fig. 2b**). Two further GA-related orthologs were identified: *RGA TARGET 1* (*RGAT1*), a direct target of GA signalling in *Arabidopsis* pollen development^38^, and a *GA 3-OXIDASE* biosynthesis gene significantly decreased in abundance (Padj < 0.05) at 130d (**Supplementary Data 2**), resembling a known homeostatic response to GA signalling^39^. These results suggested a role for GA in promoting *Ceratopteris* sporing. We verified this through exogenous application of GA3, which significantly accelerated the sporing transition (p < 0.05) (**Fig. 2c-h; Supplementary Data 1**). Our results thus support a conserved GA function between ferns and angiosperms promoting the sporophyte shoot reproductive transition.

### Sporophyll reproductive candidates include meiotic and floral development gene orthologs

Hierarchical clustering by normalised expression successfully identified differential expression of candidate genes across sporophyll development, with seven clusters exhibiting maximum abundance in different stages (**Fig. 3a**). 54.7% of candidates possessed GO term annotations, with reproduction (11.5%) and gene regulatory GO terms (6.9%) both significantly enriched (p < 0.05) (**Supplementary** Fig. 3). Reproductive GO terms related to meiosis or to reproductive shoot/organ development. Most meiosis-associated GO terms were significantly enriched in mature sporophylls (cluster 3) (**Fig. 3b**) consistent with our developmental characterisation (**Extended Data Fig. 2**), and in the sporophyll-specific repression category (**Supplementary** Fig. 4). Through these we identified orthologs of *ZIP4*, *DISRUPTED MEIOTIC CDNA1* (*DMC1*), *MSH4*, *MSH5* and a *SHUGOSHIN* (*SGO*)-like homolog (**Supplementary Data 3**), which are all conserved regulators of meiosis^40,41,42,43,44^.

**Fig. 3.**
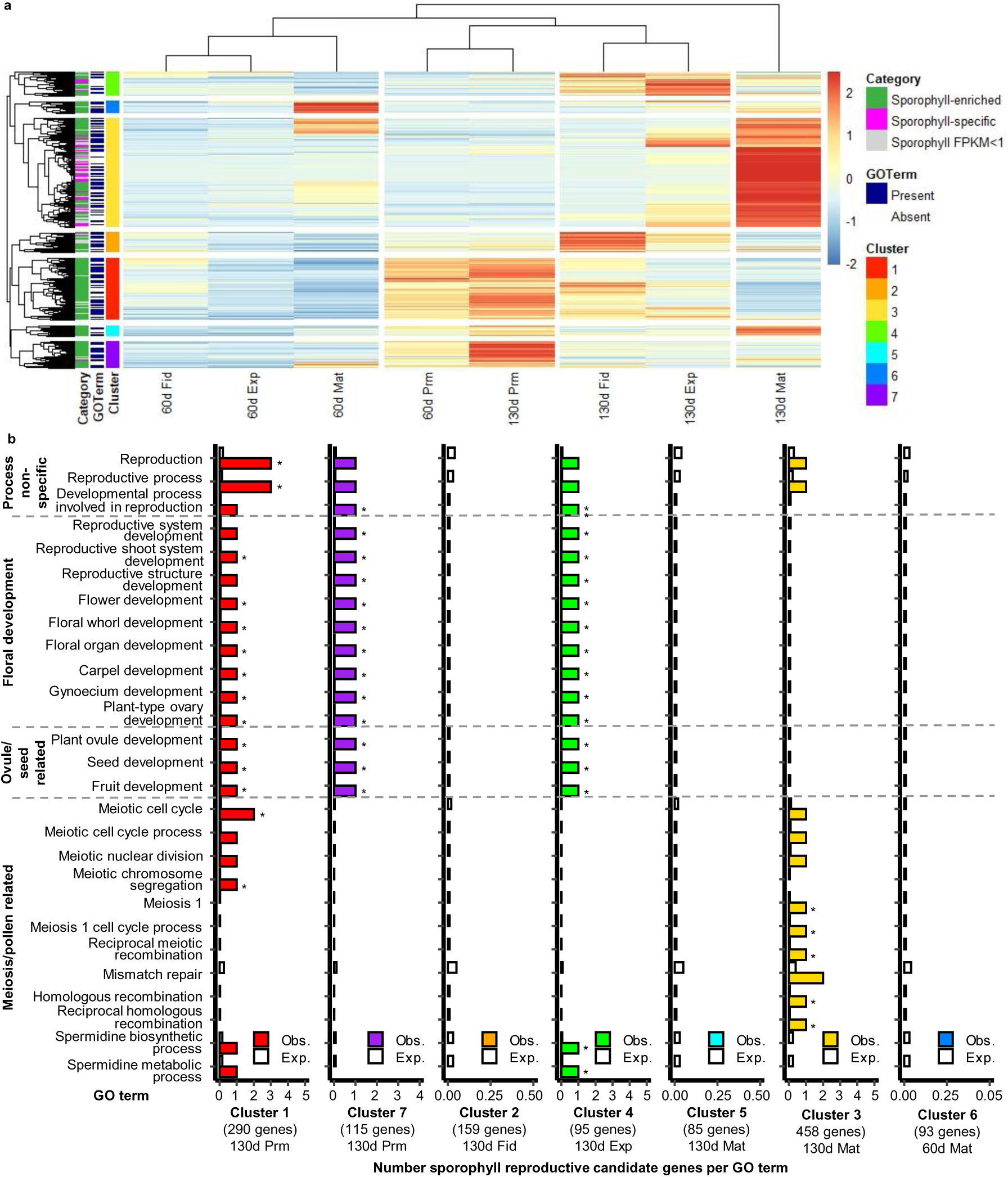
*Ceratopteris* sporop hyll candidate reproductive genes include orthologs conserved with *Arabidopsis* floral development and meiosis. **a**, Clustering sporophyll reproductive candidate genes based on normalised expression identified seven discrete clusters of genes with maximum expression in specific stages of frond development. **b**, Reproductive GO term enrichment (p < 0.05) across the sporophyll clusters identified in (**a**) in order of their peak in abundance during sporophyll development, showing the frequency of sporophyll reproductive candidate genes with enriched reproductive GO terms against the frequency expected based on the whole genome. All reproductive GO terms in which significant enrichment was detected are shown except spermine-related terms, which follow the same pattern as spermidine (**Supplementary Data 3**). Enrichment of reproductive GO terms within different expression categories are shown in **Supplementary** Fig. 4. Obs., observed frequency, Exp., expected frequency. Asterisks denote a significant increase (p < 0.05) in the frequency of genes with that GO term compared to the expected frequency.

Reproductive shoot GO terms were significantly enriched in sporophyll primordia (clusters 1 and 7) and expanding sporophylls (cluster 4) (**Fig. 3b**) and sporophyll-enriched candidates (**Supplementary** Fig. 4). All related to three MADS-box orthologs including *CMADS1* (**Supplementary Data 3**). Post-translational regulation GO terms were significantly enriched in cluster 1 (**Supplementary** Fig. 5) including an ortholog of *MERISTEMATIC RECEPTOR-LIKE KINASE* (*MRLK*), which targets floral MADS-box protein AGL24^45,46,47^, and *CrEMS1-2*^28^, an uncharacterised ortholog of *EXCESS MICROSPOROCYTES1* (*EXS/EMS1*)^48,49^ (**Supplementary Data 3**). These findings suggest conservation between the *Ceratopteris* sporophyll and *Arabidopsis* floral gene networks.

### Identifying reproductive gene networks in the *Ceratopteris* gametophyte

The *Ceratopteris* hermaphrodite gametophyte (from hereon ‘gametophyte’) develops archegonia from a ‘notch’ meristem^50,51^. To capture gene expression associated with female reproductive development, we confirmed that fertilization and embryo development could be synchronised experimentally by restricting water availability during development and then flooding once, with the majority of gametophytes bearing visible embryos 4 days after flooding (DAF) (**Extended Data Fig. 4a-i**; **Supplementary** Fig. 6**, Supplementary Data 4**). The notch meristem also exhibited fertilization-dependent shutdown by 2 DAF (**Extended Data Fig. 4j**). Genes relating to archegonium formation, fertilization and post-fertilization development were thus identified by comparing between unflooded gametophytes and 1, 2 and 4 DAF, dissecting out the notch region (‘Ntc’) (**Fig. 4a-d**) and using the surrounding thallus (‘Thl’) as a vegetative control (**Fig. 4e-h**), unavoidably including antheridia at the thallus margin (**Extended Data Fig. 4a,c,e**). The 4 DAF notch was subdivided into embryo-bearing archegonia (Aeb) and embryo-less archegonia (Arc) (**Fig. 4d**). To discriminate embryo from archegonium gene networks, we isolated 4 DAF embryos as an additional control (‘Dem’) (**Supplementary** Fig. 1).

**Fig. 4.**
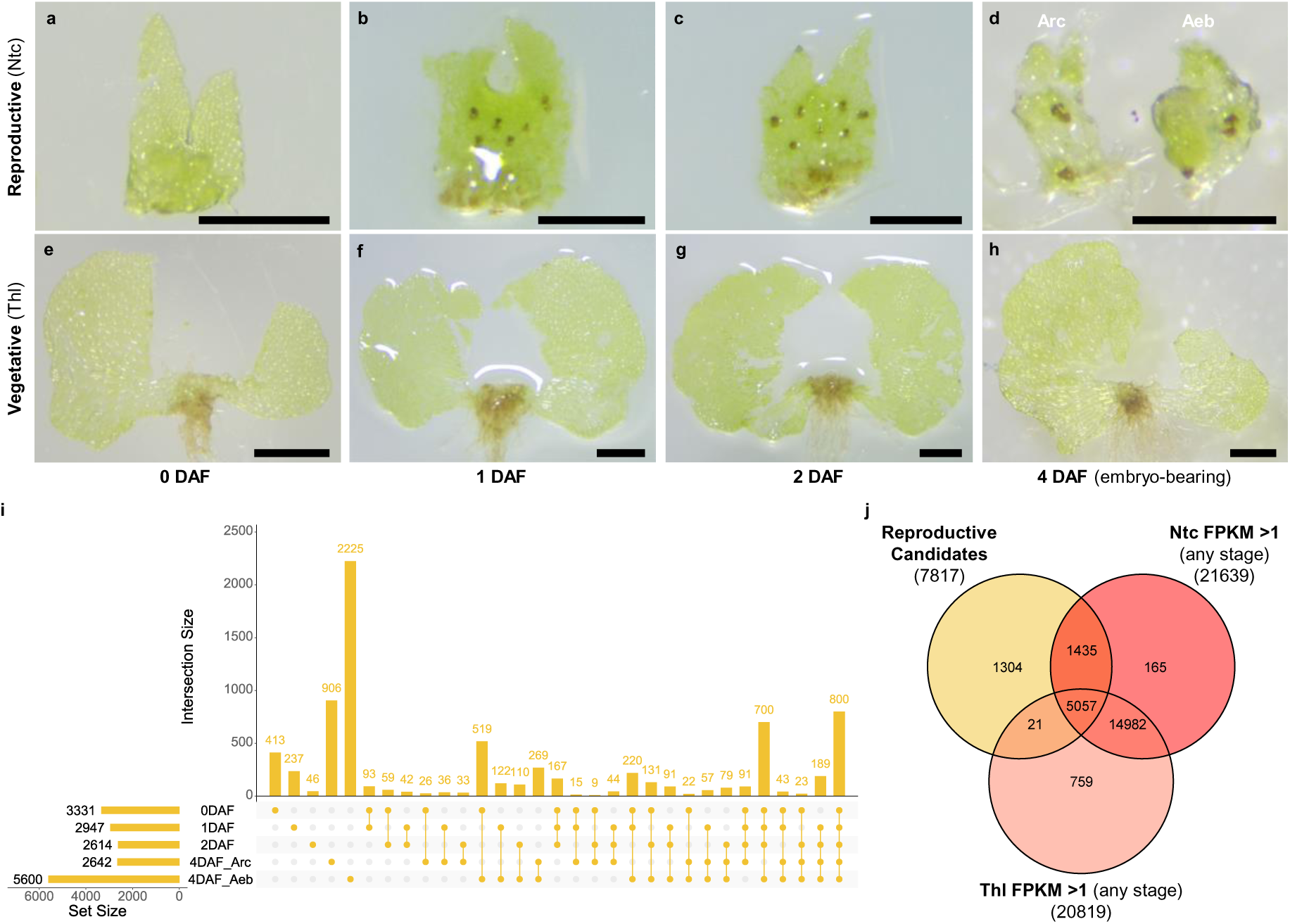
Genes associated with archegonium initiation, fertilization and post-fertilization development were identified in the *Ceratopteris* gametophyte notch. **a-h**, Reproduction-associated genes were identified by dissecting gametophytes into the notch region (Ntc), comprising the notch meristem and developing archegonia (**a**-**d**), and the surrounding thallus (Thl), containing vegetative tissue and antheridia (**e**-**h**), before fertilization (0 DAF; **a**,**e**), shortly after flooding (1 DAF and 2 DAF; **b**,**c**,**f**,**g**) and once an embryo was visible (4 DAF; **d**,**h**). 4 DAF Ntc samples were subdivided into embryo-bearing archegonia (Aeb) and embryo-less archegonia (Arc). Scale bars = 500 µm. **i**,**j**, 7187 nonredundant candidates were identified with significantly increased abundance (Padj < 0.05, log2fold expression change > 1) in Ntc vs Thl samples within at least one gametophyte developmental stage (**i**). These were categorised based on transcript abundance (**j**): candidates expressed at FPKM >1 in at least one Ntc sample but not in any Thl sample were categorised as ‘notch-specific’; those expressed at FPKM >1 in both Ntc and Thl as ‘notch-enriched’; and those with abundance of FPKM <1 in all Ntc samples as ‘notch-FPKM<1’.

PCA and correlation analysis found that biological replicates clustered more closely within each sample than between (**Extended Data Fig. 3b, Supplementary** Fig. 7) except for 1 DAF and 2 DAF (which overlapped within Ntc and Thl). PC1 separated Ntc from Thl and DAF separated progressively along PC2 (except for Dem). 7817 reproductive candidate genes were identified with increased abundance in Ntc compared to Thl (Padj < 0.05, log2foldchange >1) within at least one developmental stage (**Fig. 4i; Supplementary Data 5**). Similar to the sporophyll, candidate genes were categorised as ‘notch-specific’ (1435) and ‘notch-enriched’ (5057) (**Fig. 4j**), representing potential direct reproductive regulators and downstream developmental gene targets. 1325 remaining candidate genes were expressed at FPKM<1 in the notch.

### Notch candidates include ovule gene orthologs and regulators of post-fertilization development

52.0% of notch candidates possessed GO term annotation (**Supplementary** Fig. 3). Cell division and the cell cycle were the second and fourth most-prevalent categories of significantly-enriched GO terms (p < 0.05), reflecting the notch’s meristematic status. Fewer reproductive GO terms were enriched than in the sporophyll (1.8%) but a greater proportion related to gene regulation (13.7%), including additional post-transcriptional and epigenetic categories.

Hierarchical clustering of candidates by normalised expression successfully identified six clusters with maximum abundance at different notch developmental stages (**Fig. 5a**). Clusters 3 and 4 were significantly enriched in notch-specific candidates (Padj < 0.05) and 1, 2 and 5 in notch-enriched candidates (**Supplementary Data 6**). Reproductive GO terms were significantly enriched (p < 0.05) predominantly in clusters 2 and 3, associated with the Aeb (**Fig. 5b**). A subset were significantly enriched in transcripts with significantly greater abundance (Padj < 0.05, log2fold > 1) in Dem vs Aeb (**Fig. 5c**), including *CMADS1* (**Supplementary Data 3**). Surprisingly, many enriched reproductive terms were meiosis-related (**Fig. 5b**), identifying the same *ZIP4* and *SGO* transcripts as in the sporophyll and orthologs of *BRCA2* and *RAD51*^52,53^ (**Supplementary Data 3**). Clusters 2 and 3 were also specifically significantly enriched in post-transcriptional and epigenetic GO terms (**Extended Data Fig. 5a,b**). The associated candidate genes included orthologs of *Arabidopsis* genes from across ovule and seed development (**Extended Data Fig. 6**) and two previously-published *Ceratopteris* embryo-expressed genes (*CrANT* and *CrLFY1*)^54,55^ (**Supplementary** Fig. 8). Reproductive GO terms were not significantly enriched (p > 0.05) in pre-fertilization archegonia (cluster 4) (**Fig. 5b**), but enriched gene regulatory GO-terms identified orthologs of the meristematic genes *CLAVATA1* (*CLV1*)^56^ and *LATERAL SUPPRESSOR* (*LAS*)^57^ (**Supplementary Data 3**). Interestingly, pollen recognition GO terms linked to homologs of pollen self-recognition genes^58^ were associated with archegonia, being significantly enriched in 4 DAF embryo-less archegonia (cluster 5) (**Fig. 5b**), notch-specific candidates (**Supplementary** Fig. 9) and in genes with increased abundance in Aeb vs Dem (**Fig. 5c**). These results suggest that angiosperm reproductive networks, including seed development, are conserved with the fern archegonium.

**Fig. 5.**
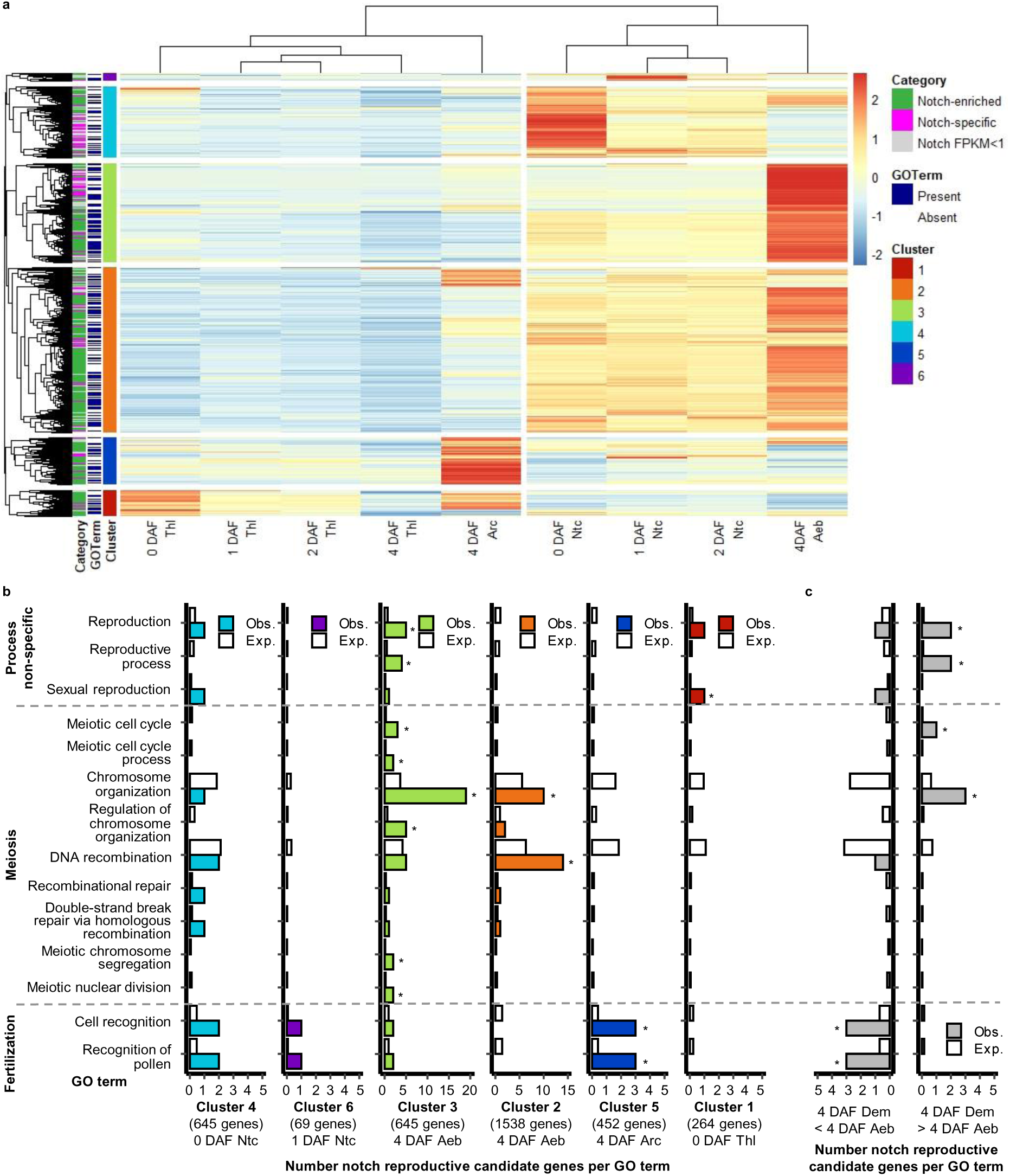
*Ceratopteris* notch candidate reproductive gene networks include conserved orthologs of meiosis regulators. **a**, Clustering of notch reproductive candidate genes based on normalised expression identified six discrete clusters of genes with maximum expression in specific tissues during gametophyte reproductive development. **b**, The frequency of observed notch reproductive candidate genes with individual reproductive GO terms against the frequency expected based on the whole genome, analysed within each cluster identified in (**a**) following the stages of notch development during fertilization (as indicated). All reproductive GO terms in which significant enrichment was detected are shown (**Supplementary Data 3**). Enrichment of reproductive GO terms within different expression categories are shown in **Supplementary** Fig. 9. Obs., observed frequency, Exp., expected frequency. Asterisks denote a significant increase (P < 0.05) in the frequency of genes with that GO term compared to the expected frequency.

Further evidence for this hypothesis emerged, with two angiosperm post-fertilization seed mechanisms identified and validated in *Ceratopteris*. Hormone response GO terms significantly enriched prior to and during fertilization (clusters 4 and 6) and in candidates with significantly greater abundance in Aeb vs Dem (**Extended Data Fig. 7a-c**) were found to relate mostly to auxin (**Supplementary Data 3**), a critical regulator of post-fertilization seed development that can drive parthenocarpic seed and fruit development^59,60,61^. To test its role in *Ceratopteris* fertilization-dependent development, exogenous auxin was applied to gametophytes at flooding. Auxin-treated gametophytes successfully bore embryos (**Extended Data Fig. 7d**), while embryo-less archegonia of the same gametophytes exhibited ectopic cell divisions not detectable in mock-treated controls (**Extended Data Fig. 7e,f**). Carbohydrate metabolism-related GO terms exhibited similar archegonium-specific enrichment (**Extended Data Fig. 8a-c**). In angiosperms, sugar signalling inhibits post-fertilization seed abortion^62,63^ and promotes apogamy in *Ceratopteris*^64^. To test the effect of carbohydrate status on *Ceratopteris* sexual reproduction, gametophytes were flooded with a 2% sucrose solution vs water. The proportion of gametophytes bearing embryos was not affected by sucrose treatment (p = 0.5393) but the mean number of embryos per embryo-bearing gametophyte significantly increased (p < 0.0001; **Extended data Fig. 8d-g**). These results suggest post-fertilization regulatory mechanisms are conserved between the fern archegonium and angiosperm seed.

### Ceratopteris sporophyll and notch candidate gene networks are conserved with distinct Arabidopsis reproductive organs

*Ceratopteris* reproductive candidates partially overlap between the sporophyll and notch (44.5% and 11.9%, respectively) (**Fig. 6a**), suggesting distinct networks. We quantified conservation between these two organs and *Arabidopsis* reproductive gene networks using gene orthology, identifying all *Arabidopsis* gene orthologs of these populations (**Fig. 6b,c; Supplementary** Fig. 10**; Supplementary Data 7**) and testing for enrichment of these within published *Arabidopsis* reproductive datasets^65,66,67,68,69,70,71,72,73,74,75,76,77^ compared to vegetative-specific genes^73^.

**Fig. 6.**
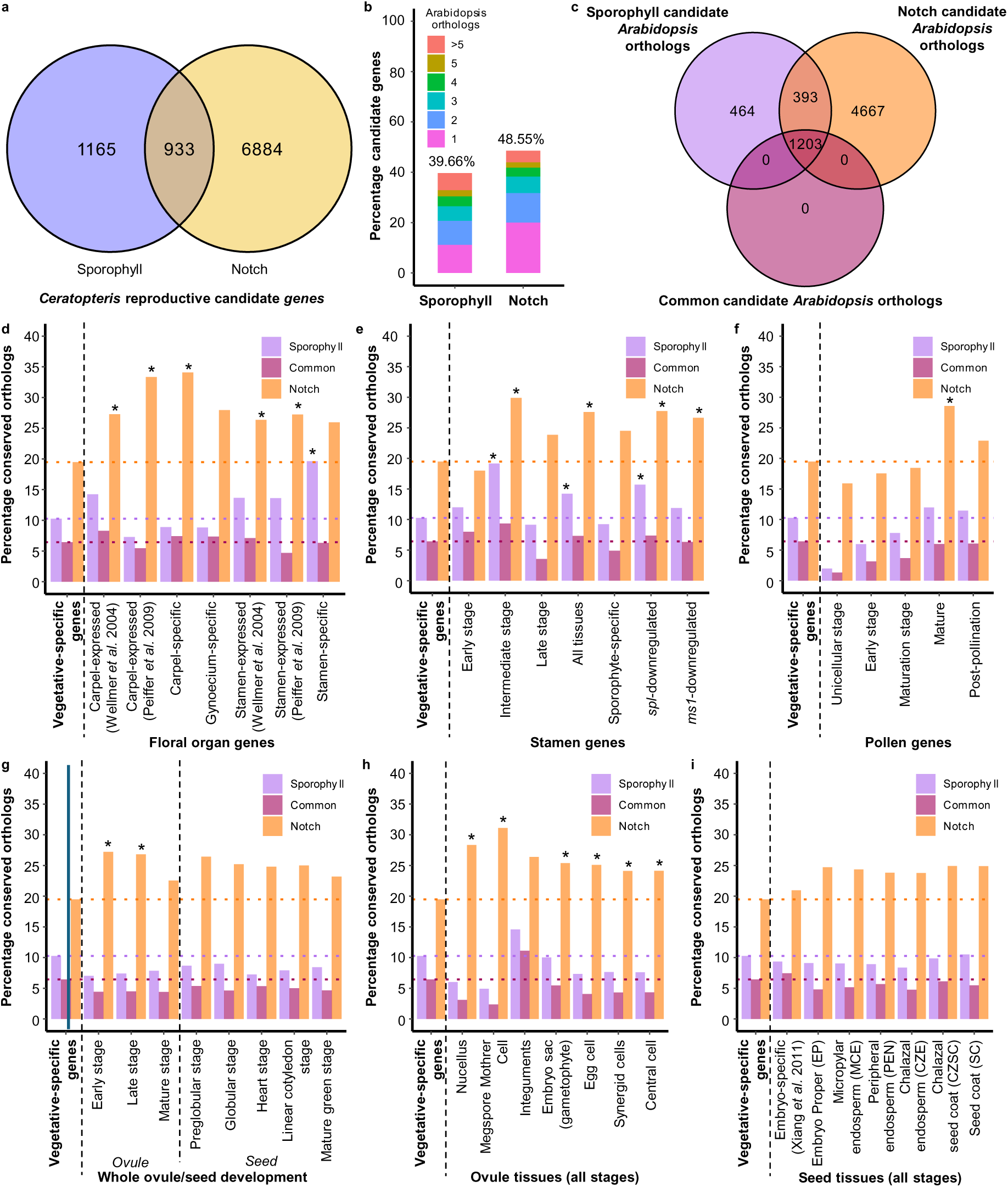
*Ceratopteris* candidate sporophyll and notch gene networks show separate conservation with distinct *Arabidopsis* reproductive organ networks. **a-c**, Sporophyll and notch reproductive candidates are partially overlapping (**a**). All direct *Arabidopsis* orthologs of these candidates were identified (**b**), of which 83.1% and 90.1% respectively possessed 5 *Arabidopsis* orthologs or less. Populations of nonredundant *Arabidopsis* orthologs for sporophyll, notch and common candidate genes were thus established (**c**) to enable enrichment testing against *Arabidopsis* reproductive gene datasets. **d-i**, Enrichment testing of *Arabidopsis* gene orthologs for the sporophyll, notch and common orthologs within published datasets of *Arabidopsis* reproductive genes expressed in floral organs^67,71^ (**d**), stamen development^69^ (**e**), pollen development^65,66^ (**f**), whole ovule/seed development^68,73,77^ (**g**), ovule tissues^70,72,74,75^ (**h**) and seed tissues^76,77^ (**i**) vs published vegetative-specific genes^73^. Barplots show the overlap of candidate orthologs against each target as a proportion of that dataset. Asterisks denote a significant increase (Padj < 0.05) in the proportion of overlapping genes between candidate orthologs and the target reproductive organ compared to the overlap with vegetative-specific genes. Dotted horizontal lines track the baseline overlap of each candidate ortholog dataset with the vegetative control. p-values for individual comparisons are provided (**Supplementary Data 8**).

Sporophyll orthologs were significantly enriched (Padj < 0.05) only in stamen datasets (**Fig. 6d-i**), including *SPOROCYTELESS* (*SPL*)-dependent genes relating to sporogenesis^78,79^. This persisted in sporophyll-enriched orthologs only (**Supplementary Data 8**). Notch orthologs were significantly enriched in stamen, pollen, carpel and ovule datasets (**Fig. 6d-i**), with orthologs of notch-enriched candidates showing additional enrichment in post-fertilization seeds (**Extended Data Fig. 9a-c**). Notch orthologs were significantly enriched in all ovule tissue datasets tested except the integument (**Fig. 6h**). Common orthologs were not significantly enriched (Padj > 0.05) in any dataset (**Supplementary Data 8**), supporting separate conservation of gene networks between the sporophyll and the stamen, and between the notch and the stamen, pollen and ovule/seed.

Notch-enriched orthologs significantly-enriched within the ovule formed two highly-correlated groups, between sporophytic and gametophytic tissues (**Extended Data Fig. 9d; Supplementary** Fig. 11). However, most tissue-specific and commonly-expressed subsets within and between these two groups remained significantly enriched in notch-enriched orthologs (**Extended Data Fig. 9e-g**). 43.7% of orthologs significantly-enriched in the seed were not present in the ovule (**Supplementary** Fig. 11). Notch-enriched orthologs were more equally correlated between seed tissues and commonly-expressed subsets remained significantly enriched (**Extended Data Fig. 9h,i**). However, we found that seed-specific genes were not significantly enriched in notch-enriched orthologs (Padj > 0.05) (**Extended Data Fig. 9j**), suggesting a common evolutionary origin for ovule and seed gene networks from the notch.

To determine if specific stages of *Ceratopteris* sporophyll and notch development were conserved with *Arabidopsis* reproductive organs, reciprocal enrichment testing was undertaken using all *Ceratopteris* reproductive genes orthologous to the conserved genes significantly-enriched within each *Arabidopsis* organ (**Extended Data Fig. 10a,b; Supplementary Data 9**) within sporophyll and notch expression clusters. Sporophyll orthologs of conserved stamen genes were not significantly enriched in any cluster compared to all sporophyll candidates (Padj > 0.05; **Extended Data Fig. 10c,d**). Notch orthologs of conserved stamen genes were significantly enriched (Padj < 0.05) (**Extended Data Fig. 10e**), which could be ascribed to conserved pollen genes. Conserved ovule genes were significantly enriched in the 4 DAF Aeb (clusters 2 and 3), which could be ascribed to genes conserved with the nucellus, MMC and embryo sac central cell (**Extended Data Fig. 10f-i**). To test the developmental origin of post-fertilization seed gene networks, enrichment testing was repeated using orthologs of notch-enriched genes conserved in *Arabidopsis* (**Supplementary** Fig. 12). Conserved embryo, endosperm and seed coat orthologs were significantly enriched in cluster 3, further supporting the conservation of gene networks between the ovule/seed and post-fertilization archegonium.

## DISCUSSION

Our mRNA-seq analysis identified gene conservation at points throughout *Ceratopteris* and *Arabidopsis* reproductive development, supporting the hypothesis that seed plant reproductive innovations arose by repurposing ancestral reproductive gene networks. This allows us to provisionally assign development origins to these networks.

GA signalling was identified and validated as promoting the *Ceratopteris* sporing transition. GA signalling promotes the *Arabidopsis* floral transition by indirectly upregulating floral meristem identity genes^80^ including MADS-box genes^81^. MADS-box gene clades regulating floral meristem and floral organ identity are angiosperm or seed-plant specific^4,82^ but *Ceratopteris* homologs were identified with expression patterns consistent with downstream targets of the sporing transition. This suggests that the GA module of angiosperm shoots was co-opted from an ancestral mechanism in the sporing transition.

Angiosperm MADS-box proteins specify different reproductive organ identities during flower development, with similar functions proposed in gymnosperms^25^. We further identified MADS-box genes (including *CMADS1*) in early sporophyll development, together with an upstream floral regulator of MADS-box activity (*MRLK*). These were sporophyll-enriched rather than specific, and previously-published *CMADS1 in situ* localisation data^24^ supports dual vegetative/reproductive expression. Sporophyll candidates were also significantly enriched in stamen genes, consistent with the *Adiantum capillus-veneris* homosporangium^30^. Floral-specific networks may thus have arisen from an ancestral sporophyll programme.

We identified orthologs of genes regulating male and female meiosis within sporophyll candidates^40,41,42,43^ but two of these and further meiosis genes (*BRCA2*, *RAD51*)^52,83^ with highly-conserved functions^84,85,86,87^ were also identified in the post-meiotic gametophyte notch. *AtBRCA2A* and *-B* have a dual somatic function^83^, likewise paralogs of *AtRAD51* (*AtRAD51C*)^88^. Their presence in the actively-dividing notch could reflect a similar role in *Ceratopteris*. Notch candidates also included orthologs of female-specific meiosis regulators^89^: *AtDRM1* and *-2* prevent the formation of ectopic MMCs^90^. These results suggest that homosporous meiosis genetically resembles seed plant microsporogenesis, consistent with their morphological similarity^19^, while female meiosis could have diverged by co-opting notch gene networks already utilising meiosis genes.

Contrary to expectations, *Arabidopsis* ovule/seed genes were conserved specifically with the *Ceratopteris* post-fertilization archegonium. This raises the intriguing hypothesis that the diploid ovule/seed body-plan arose by co-opting gene networks from archegonium development. Although counterintuitive, this accords with the striking functional and developmental similarities between the post-fertilization fern archegonium and the seed^11^, together with the conservation of carbohydrate status and auxin regulating archegonium and seed post-fertilization development. The feasibility of mis-expressing developmental programmes between gametophyte and sporophyte generations has been demonstrated in *Physcomitrium patens*, where loss of *CURLY LEAF* caused activation of sporophyte development in the gametophyte^91^. It is therefore possible that mis-activation of the post-fertilization archegonium gene network during sporophyll development contributed to the formation of the ovule.

## ONLINE METHODS

### Plant material and growth conditions

*Ceratopteris richardii* lab strain Hn-n^92^ was used for all experiments. Spore sterilization, plating and gametophyte growth on C-fern 1% agar medium (pH6.0) under tissue culture conditions were performed as previously described. Plates were kept humid by adding 1.5 ml sterile water at 3 and 7 days after sowing, with no water added beyond this time to prevent premature sperm release and fertilization. Fertilization was triggered at 14 days by flooding plates with 7.5 ml sterile water, with embryo development assumed to have begun on the day of flooding. individual sporophytes transferred individually to fresh C-fern agar media once visible (9-11 days after flooding [DAF]). Individual sporophytes were transferred to sterile GA-7 Magenta vessels containing 100 ml C-fern 1% agar medium (pH6.0) when 2-3 fronds and elongating roots were visible (14-16 DAF) and wetted with 5 ml sterile water. Sporophytes were subsequently maintained in Magenta vessels until 130 days old (**Supplementary** Fig. 13), with sterile water re-applied whenever no surface water was visible (typically every 2-3 weeks). At 100 and 120 days 5 ml of sterile C-fern liquid media (pH6.0) was applied instead of water to maintain healthy growth. Sporophytes grown for harvesting 25 day-old samples were transferred to 90 mm plates at a density of 30-40 per plate and grown there until harvested. All plant material was grown in an MLR-352H-PE controlled environment cabinet (PHC Europe, Breda, Netherlands) at 28°C, constant 50% relative humidity and a 16hr light/ 8hr dark photoperiod at a fluence of 150 μmol m^-2^ s^-1^.

### Embryo development synchronisation

To confirm the efficacy of restricted watering at synchronising embryo development, populations of gametophytes were grown at a density of 200-300 spores per plate under tissue culture conditions, with parallel growth regimes applied either restricted watering (as described above) or ‘well-watered’ conditions with sterile water applied at 9 and 11 DAS to maintain a constant layer of surface water, at ratio of 2 restricted water:1 well-watered. At 14 DAS, all well-watered plates and half of the restricted water plates were flooded with sterile water as described above (‘+Flooding’), with the remaining restricted water plates left dry (‘-Flooding’). One plate per treatment was sacrificed for analysis at 14 DAS (Restricted-water, Well-watered), 4 DAF and 7 DAF (Restricted water -Flooding, Restricted water +Flooding, Well-watered +Flooding). From each plate 50 hermaphrodite gametophytes were selected at random and scored for the presence of an embryo using an ultraZOOM-2 stereomicroscope (GT Vision Ltd, Newmarket, UK) at 20x magnification. The experiment was replicated on four different occasions.

To characterise the timing of post-fertilization embryo development and archegonium production in response to flooding, populations of gametophytes were grown at a density of 200-300 spores per plate under restricted water conditions (as described above) and flooded at 14 DAS. At 0 (unflooded), 1, 2, 3, 4, and 7 DAF single plates were sacrificed for analysis. From each plate 50 hermaphrodite gametophytes were selected at random and scored for the presence of an embryo (as described above). At 1-7 DAF gametophyte selection was restricted to those with evidence of fertilization in response to flooding (brown archegonia). The experiment was replicated on five different occasions. During three of these replicates the number of archegonia per gametophyte were counted alongside scoring for the presence of an embryo. Numbers of embryo-bearing and embryo-less flooded gametophytes recorded per replicate varied due to availability within the population.

### Sporing transition characterisation

Sporophytes were grown in Magenta vessels (as described above) in populations of 25. Eleven replicate populations were grown, with populations initiated at intervals over 13 months. From 4 weeks old onwards, the number of sporophytes with visible sporophylls within each population was recorded once per week ±1 days until 19 weeks old, by which time 100% of sporophytes had begun to spore. For nine of these replicates the number of fronds per sporophyte was also recorded weekly (**Supplementary Data 1**).

### Sporangium development characterisation

Single sporophylls were collected from 130d sporophytes, secured dorsal-side down to microscope slides using double-sided tape and pinnae dissected open under a stereomicroscope using hypodermic needles as micro-scalpels to expose developing sporangia on the ventral surface. Sections of pinnae were recorded photographically to capture sporangium planar area. Sporangia were recorded at two locations per pinna (base and tip) and from two pinnae per frond, positioned basally and distally. Four sporophytes were sampled per frond developmental stage. Sporangia areas were measured from scaled micrographs using the FIJI image analysis software package^94^.

### mRNA-seq sample preparation

Tissue samples for mRNA-seq were harvested by dissection and immediately frozen in liquid nitrogen. Sequential stages of sporophyte frond development were harvested within 2 hours per population grown, with replicate populations harvested between 5 and 10 hours after dawn (ZT5-ZT10). Frond developmental stages were determined by eye based on frond morphology. Dissection of Apx and Prm samples was performed under a stereomicroscope using watchmaker’s forceps. Tissues were frozen immediately after dissection by collection into tubes floating on liquid nitrogen. All biological replicates comprised tissue from at least 4 independent sporophytes (**Supplementary Data 10**), harvested from one (all Mat, Exp, Fid, 60d Apx and Prm), 2-3 (130d Apx and Prm) or 3-4 independently-harvested populations (25d Apx and Prm), respectively. Exp samples were designed to encompass ‘early’ (unfurling), ‘mid’ (unfurled to 50% final size) and ‘late’ frond expansion (50-75% final size) (**Extended Data Figure 2**) within each biological replicate. Samples across frond development comprised tissue from the same harvested population(s) per biological replicate.

Gametophyte stages were necessarily harvested on separate days, individual populations harvested within 2-4 hours with replicate populations harvested between 5 and 10 hours after dawn (ZT5-ZT10). Dissection of the notch was performed under a stereomicroscope using hypodermic needles as micro-scalpels, the developmental status of each individual confirmed by eye. Each biological replicate comprised notch or thallus tissue from 150-225 individual gametophytes (**Supplementary Data 10**) pooled from 1 (0 DAF, 1 DAF, 2DAF), 2-3 (4 DAF) or 3-4 independently-harvested populations (Dem, 761-788 embryos per sample) respectively. Notch and thallus from the same individuals remained paired when preparing biological replicates for sequencing.

RNA was extracted using the Qiagen RNeasy plant mini kit with on-column DNase treatment (QIAGEN, Hilden, Germany). RNA quality was determined using a 2100 Bioanalyser RNA 6000 nano assay (Agilent Technologies, Santa Clara, USA). Individual sample RIN values are supplied in **Supplementary Data 10**. Library preparation, 150bp paired-end Illumina sequencing was performed separately for sporophyte and gametophyte samples by Novogene (Novogene UK, Cambridge UK) to a depth of ≥ 20 million reads per sample. QC data is provided in **Supplementary Data 10**.

### Imaging

Whole sporophytes and gametophyte plates were photographed using a Cybershot DSC-HXTV digital camera (Sony Corporation, Tokyo, Japan). Individual gametophytes and dissected tissue samples were photographed using an ultraZOOM-2 stereomicroscope with an attached GXCAM-U3PRO-X digital camera and GXCam software (GT Vision Ltd, Newmarket, UK). Photographs were minimally processed by adjusting brightness and contrast and sharpness for whole images.

### Data analysis

Statistical analyses of characterization experiments were conducted in R version 4.2.2^95^ using the car package^96^. Pairwise comparisons were performed using two-tailed students T-tests or Mann-Whitney nonparametric tests (indicated within each figure), following analysis of the data structure using Shapiro-Wilk for Normal distribution^97^ and Levene’s Test for equal variance^98^. Where necessary data was mathematically transformed to meet the assumptions of the statistical tests (indicated within each figure). A post-hoc correction for multiple comparisons was applied to p-values using the Benjamini-Hochberg method^99^.

Transcript assembly against the *Ceratopteris richardii* genome v2.1^32^ using HISAT2^100^ was performed by Novogene, as was annotation and quantification of transcripts and pairwise DE analysis (**Supplementary Data 2; Supplementary Data 5**). Orthology of *Ceratopteris* genes to *Arabidopsis* was identified using Orthofinder^101^ with default settings against the following proteomes: *Arabidopsis thaliana* (TAIR10)^102^, *Amborella trichopoda* (v1.0)^103^, Glycine max (Wm82.a4.v1)^104^, *Manihot esculenta*(v8.1) (https://phytozome-next.jgi.doe.gov/info/Mesculenta_v8_1), *Micromonas pusilla* CCMP1545 (v3.0)^105^), *Ostreococcus lucimarinus* (v2.0)^106^, *Oryza sativa* (v7.0)^107^, *Physcomitrium patens* (v3.3)^108^, *Quercus robor* (PM1N)^109^, *Solanum lycopersicum* (ITAG4.0)^110^, *Selaginella moellendorffii* (v1.0)^111^ *Spirodela polyrhiza* (v2)^112^ and *Volvox carteri* (v2.1)^113^. Additional GO term annotations were assigned to *Ceratopteris* genes from gene orthologs in the model species *Arabidopsis*, *Oryza*, *Selaginella, Physcomitrium* and *Marchantia*. Higher-order Venn comparisons were performed in InteractiVenn^114^). Further analysis of transcriptome data was performed in R, with cluster analysis using the pheatmap package^115^ and GO term enrichment testing using the TopGO package^116^, employing Fisher’s Exact Test. *Arabidopsis* reproductive gene datasets were obtained from past publications and filtered against published vegetative-specific genes (**Supplementary Data 8**). Enrichment testing of *Arabidopsis* orthologs of *Ceratopteris* genes against these datasets was performed in R using Fisher’s Exact Test (one tailed, greater than). The same methodology was used for reciprocal enrichment testing of *Ceratopteris* orthologs of conserved *Arabidopsis* genes within candidate expression clusters. All other plots were prepared in R using the ggplot2^117^, ggvenn^118^ and ggcorrplot^119^ packages.

## DATA AVAILABILITY

Raw sequencing read data is available via the NCBI Gene Expression Omnibus (GEO) data repository (reference GSE291236). All other raw and processed data from the experiments presented here are provided in online supplemental information.

## ACKNOWLEDGEMENTS

This research was funded by Royal Society University Research Fellowship URF\R1\191326. We are grateful to Dr Juliet Coates and Prof Daniel Gibbs for helpful discussions about mRNA-seq experimental design, Drs Megan McDonald and Peter Keane for helpful discussions about data analysis and Profs Daniel Gibbs, Christine Foyer and Eugenio Sanchez-Moran for helpful discussions about the manuscript.

## AUTHOR CONTRIBUTIONS

ARGP conceived and planned experiments, performed developmental characterisation, tissue harvesting, RNA extraction, GO term analysis and enrichment testing. MC performed gene orthology assignment. ARGP and MC wrote the manuscript.

## SUPPLEMENTARY INFORMATION

**Supplementary** Fig. 1. Additional mRNA-seq samples and developmental ranges.

**Supplementary** Fig. 2. Correlation analysis between all sporophyte mRNA-seq samples.

**Supplementary** Fig. 3. Functional categorisation of enriched GO terms in sporophyll and notch reproductive candidate genes.

**Supplementary** Fig. 4. Distribution of enriched reproductive GO terms across sporophyll reproductive candidate expression categories.

**Supplementary** Fig. 5. Distribution of enriched gene regulatory GO terms across sporophyll reproductive candidate expression categories/clusters.

**Supplementary** Fig. 6. *Ceratopteris* embryo development is synchronised through restricted watering.

**Supplementary** Fig. 7. Correlation analysis between all gametophyte mRNA-seq samples. **Supplementary** Fig. 8. Validation of notch reproductive candidate genes as previously-published embryo-related genes.

**Supplementary** Fig. 9. Distribution of enriched reproductive GO terms across notch reproductive candidate expression categories.

**Supplementary** Fig. 10. *Arabidopsis* gene orthology to *Ceratopteris* reproductive candidates by expression category.

**Supplementary** Fig. 11. Comparison of significantly enriched notch-enriched *Arabidopsis* orthologs between ovule and seed tissues.

**Supplementary** Fig. 12. Reciprocal enrichment testing of *Ceratopteris* notch-enriched orthologs of conserved *Arabidopsis* genes within notch-enriched candidates.

**Supplementary** Fig. 13. Growth of the *Ceratopteris* sporophyte under tissue culture conditions.

**Supplementary Data 1.** Sporophyte phenotypic characterisation data and statistical analysis.

**Supplementary Data 2.** Sporophyll reproductive candidate gene DE analysis.

**Supplementary Data 3.** GO term enrichment analysis of *Ceratopteris* reproductive candidate genes (sporophyll and notch)

**Supplementary Data 4.** Gametophyte fertilization characterisation data and statistical analysis.

**Supplementary Data 5.** Notch reproductive candidate gene DE analysis.

**Supplementary Data 6.** Pheatmap gene identities and additional statistical analysis

**Supplementary Data 7.** *Ceratopteris* gene orthology source data.

**Supplementary Data 8.** *Arabidopsis* gene ortholog enrichment testing source data and statistical analysis.

**Supplementary Data 9.** Conserved *Ceratopteris* gene ortholog enrichment testing source data and statistical analysis.

**Supplementary Data 10.** *Ceratopteris m*RNA-seq sample information and QC data

---

**Extended Data Fig. 1.**
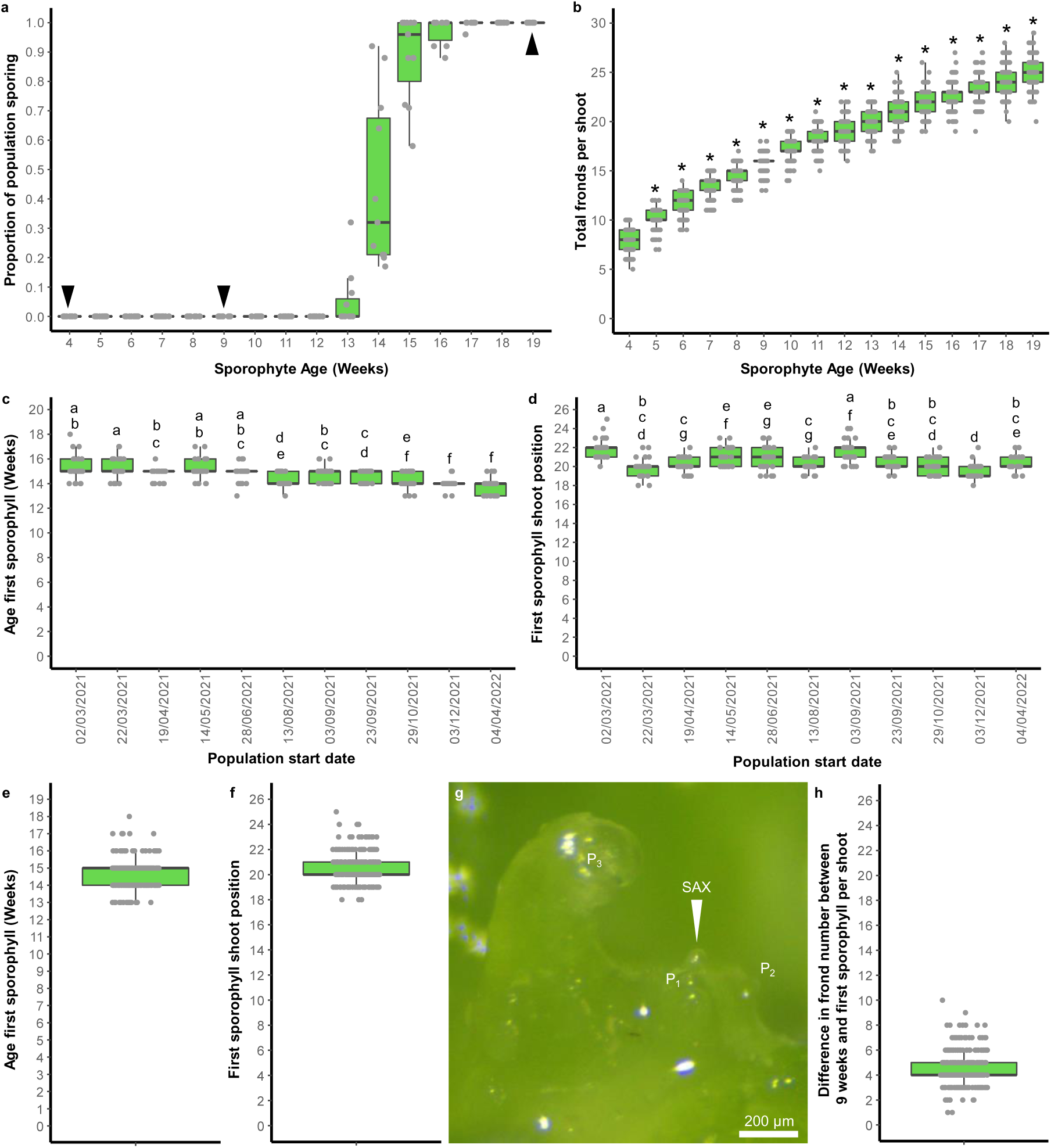
Characterising the timing and stability of sporing in the *Ceratopteris* sporophyte shoot. **a,b**, Plots show the proportion of sporophytes sporing over time (**a**) and the total number of fronds per sporophyte over the same period (**b**), confirming that shoot development remained active throughout. Asterisks in (**b**) denote a significant difference (Padj < 0.05) between the age indicated and the preceding week. **c-f**, Earliest sporing time by chronological age and shoot position, within each population grown (**c,d**) and across all sporophytes (**e,f**). Black arrowheads in (**a**) indicate shoot ages sampled for mRNA-seq. Letters in (**c**,**d**) denote statistical similarity, with different letters indicating a significant difference (Padj < 0.05) between experimental populations. **g**,**h** The 9 week-old shoot apical region contains the shoot apex (SAX) and sequentially-developing primordia (P1-P3) not visible to the naked eye and so not included in frond counts. Comparing the number of visible fronds at 9 weeks old and the first sporophyll shoot position (**h**) found a mean difference significantly greater than 3 (p < 0.0001, one-tailed Mann-Whitney test), indicating that 9 week-old shoots have not transitioned. Boxplots denote the data median and interquartile range (box) ±1.5x the interquartile range (whiskers). *n* = 11 (**a**), 225 (**b**) 25 (**c**,**d**), 271 (**e,f,h**). Pairwise comparisons were performed using two-tailed Mann-Whitney tests on square-root (**b**) and log10-transformed data (**c**,**d**), respectively (**Supplementary Data 1**).

**Extended Data Fig. 2.**
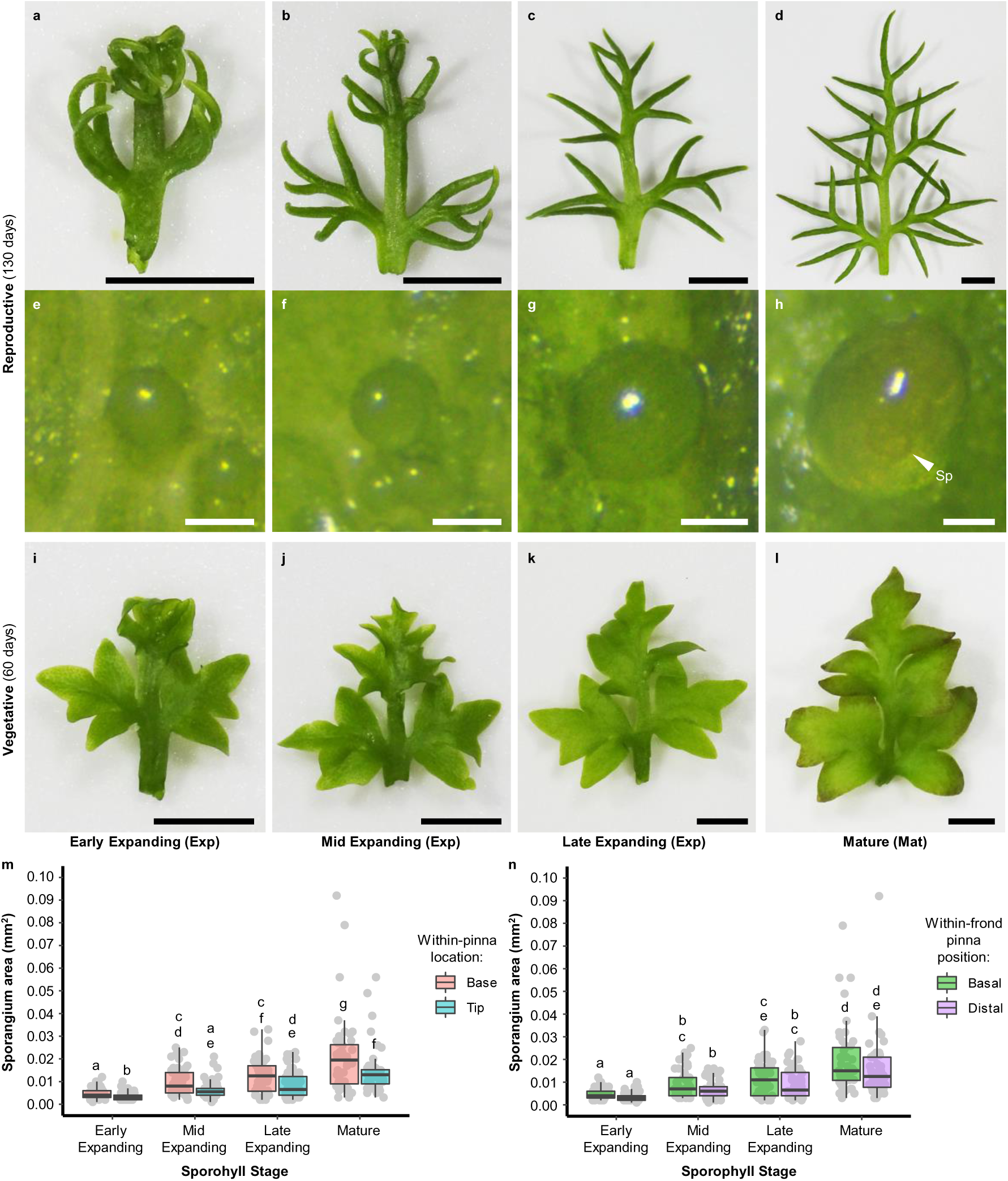
Characterisation of *Ceratopteris* sporangium development during sporophyll expansion. **a-h**, Sub-stages within ‘expanding sporophyll’ (Exp) and representative sporangia, based on frond size: ‘early expanding’ (**a**,**e**, where fronds are still unfurling); ‘mid expanding’ (**b**,**f**, from newly-unfurled fronds to approximately 50% of final size); ‘late expanding’ (**c**,**g**, from approximately 50% to 75% of final size) and ‘mature’ (**d**,**h**) where fronds are of similar size to those immediately preceding it. Post-meiotic sporangia containing individual spores (Sp) were visible in mature frond samples only. **i-l**, Expanding vegetative fronds from 60 day-old sporophytes, following the sub-stages established for sporophylls. Scale bars = 5 mm (black) and 100 µm (white). **m,n**, Sporangium planar area measured across sub-stages of frond expansion, comparing sizes within individual pinnae (**m**) and between pinnae within an individual frond (**n**). *n* =24 (4 sporangia per pinna location, measuring 6 independent sporophytes). Boxplots denote the data median and interquartile range (box) ±1.5x the interquartile range (whiskers). Letters denote statistical similarity, with different letters indicating a significant difference (Padj < 0.05) between groups. Sporangium sizes were found to overlap between adjacent frond substages. Pairwise comparisons were performed using two-tailed Mann-Whitney tests on log10-transformed data (**Supplementary Data 1**).

**Extended Data Fig. 3.**
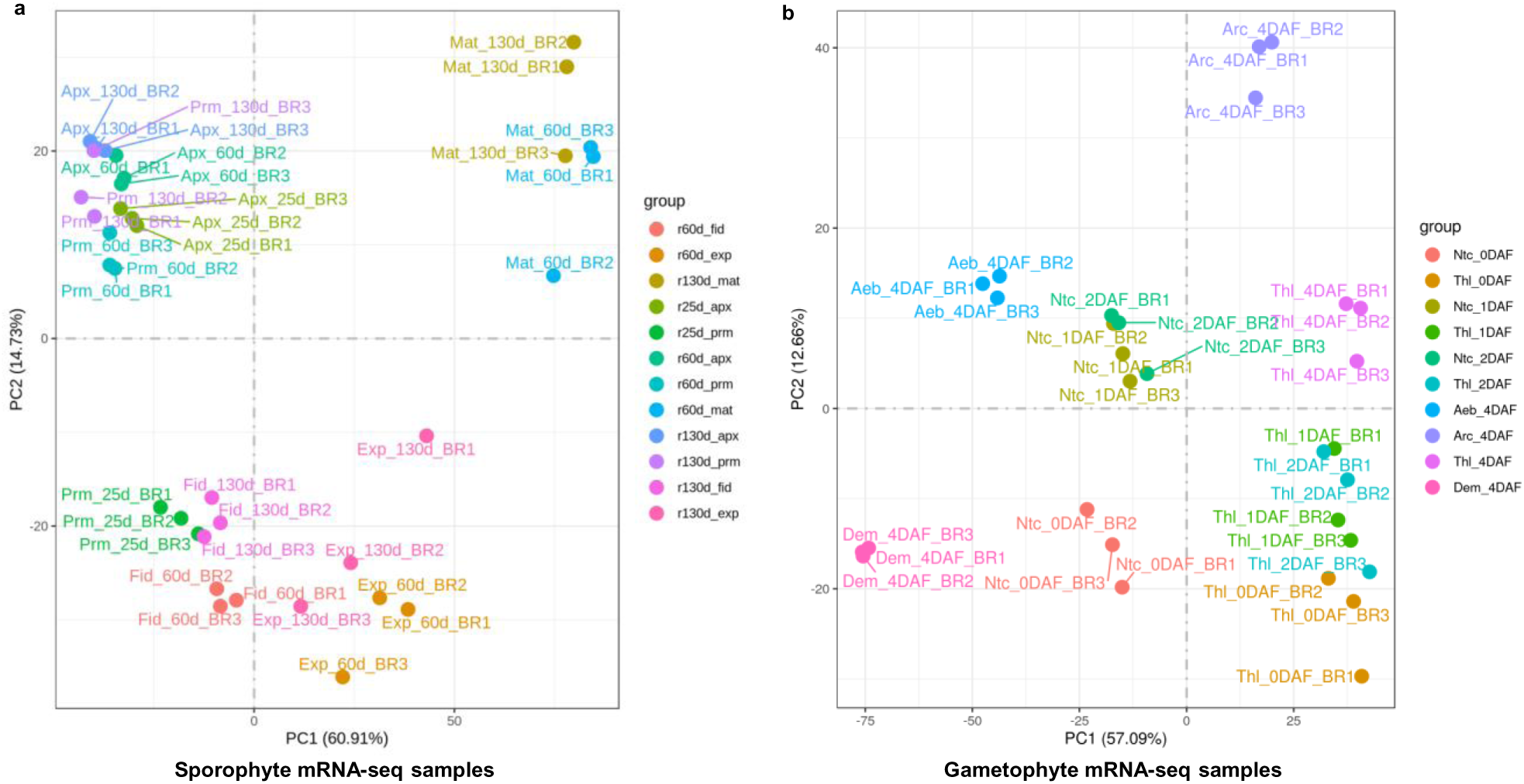
Principal Component Analysis of *Ceratopteris* mRNA-seq data. **a**, Biological replicates within sporophyte samples typically cluster more closely together than with replicates of other samples, the exceptions being primordia samples from 25d sporophytes (Prm_25d) which cluster more closely with fiddleheads. PC1 separates frond developmental stages progressively in order of developmental stage. PC2 separates vegetative and reproductive samples within each frond stage. **b**, Biological replicates within gametophyte samples typically cluster more closely together than with replicates of other samples, with the exception of 1DAF vs 2DAF within notch and thallus samples. PC1 separates notch and embryo-bearing samples from thallus samples, with embryo-less 4DAF notch tissues (Arc_4DAF) falling between notch and thallus samples on this axis. PC2 separates samples progressively by age (with 1DAF and 2DAF overlapping), with the exception of dissected embryos (Dem_4DAF).

**Extended Data Fig. 4.**
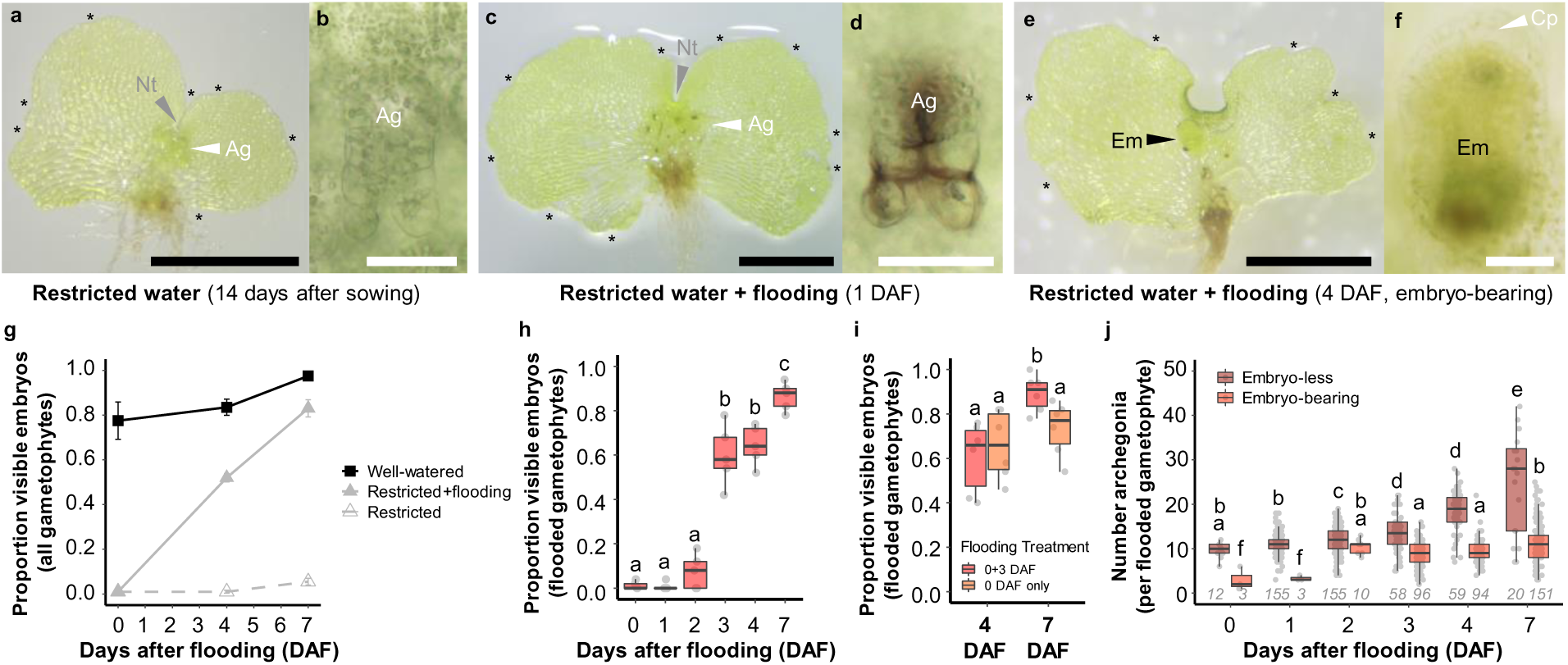
Restricted watering synchronises *Ceratopteris* fertilization. **a-f**, When grown with restricted water, mature unfertilized gametophytes (**a**) have green archegonia (Ag) (**b**) arising sequentially from the notch meristem (Nt), while antheridia (asterisks) develop at the thallus margin. By 1 day after flooding (DAF) (**c**) all archegonia turn brown (**d**). Typically a single embryo (Em) develops per gametophyte (**e**), the enclosing archegonium expanding into a calyptra (Cp) (**f**). Scale bars = 1 mm (black), 50 µm (white). **g**, Proportion of gametophytes (mean ± SE) with visible embryos under well-watered conditions or if watering was stopped at 7 days after spore sowing (‘Restricted’). By 14 days after sowing (0 DAF) 77.5 ± 12.5% of well-watered controls bore embryos at various developmental stages (**Supplementary** Fig. 6) compared to 1.0 ± 1.0% under restricted water. Fertilization could be triggered in restricted-water populations by flooding, with 83.0 ± 5.74% bearing visible embryos by 7 DAF compared to 5.5 ± 2.2% without flooding. *n* = 4. **h**-**i**, Proportion of flooded gametophytes with visible embryos per DAF. Most embryos were first visible by 3-4 DAF (**h**). If re-flooded subsequently (‘0+3 DAF’) this proportion increased between 4 and 7 DAF (p < 0.05) (**h,i**), suggesting additional fertilization events, but if flooded only once (‘0 DAF only’) the proportion did not increase beyond 4 DAF. *n* = 5. **i**, Number of archegonia per flooded gametophyte per DAF, comparing between gametophytes with visible embryos (‘embryo-bearing) and those without (‘embryo-less’). Archegonium numbers stopped increasing in embryo-bearing gametophytes after 2 DAF but continued in embryo-less individuals. *n* shown in grey italic for each embryo status vs DAF combination. Boxplots show the median and interquartile range (box) ±1.5x interquartile range (whiskers). Letters denote statistical similarity, with different letters indicating a significant difference (Padj < 0.05) between groups. Pairwise comparisons were performed using two-tailed Mann-Whitney tests on logit (**h**,**i**) or log10-transformed (**j**) data (**Supplementary Data 4**).

**Extended Data Fig. 5.**
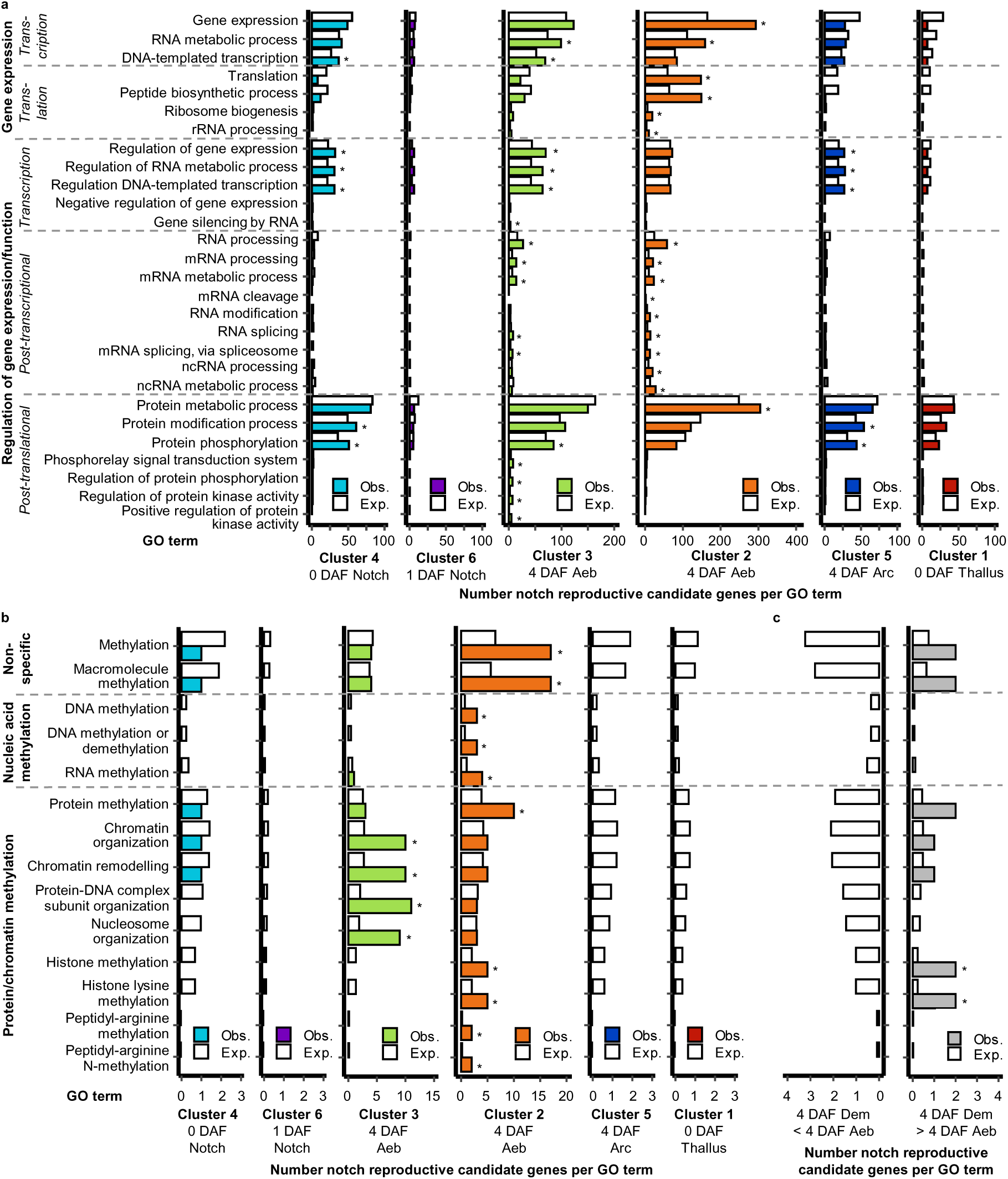
Post-transcriptional gene regulation and epigenetic GO terms are enriched specifically in the *Ceratopteris* embryo-bearing archegonium. **a-c**, Enrichment of GO terms relating to gene expression (**a**) or epigenetic regulation (**b**) within each expression cluster following the stages of notch development (see Fig. 5), and epigenetic GO terms enriched within genes showing significantly increased or decreased abundance in 4 DAF embryo-bearing archegonia vs 4 DAF dissected embryos (**c**). Bar plots show the observed (‘Obs.’) and expected (‘Exp.’) gene frequencies per GO term. Asterisks denote a significant increase (p < 0.05) in genes with that GO term compared to that expected based on the whole genome. Individual p-values are provided (**Supplementary Dataset 3**).

**Extended Data Fig. 6.**
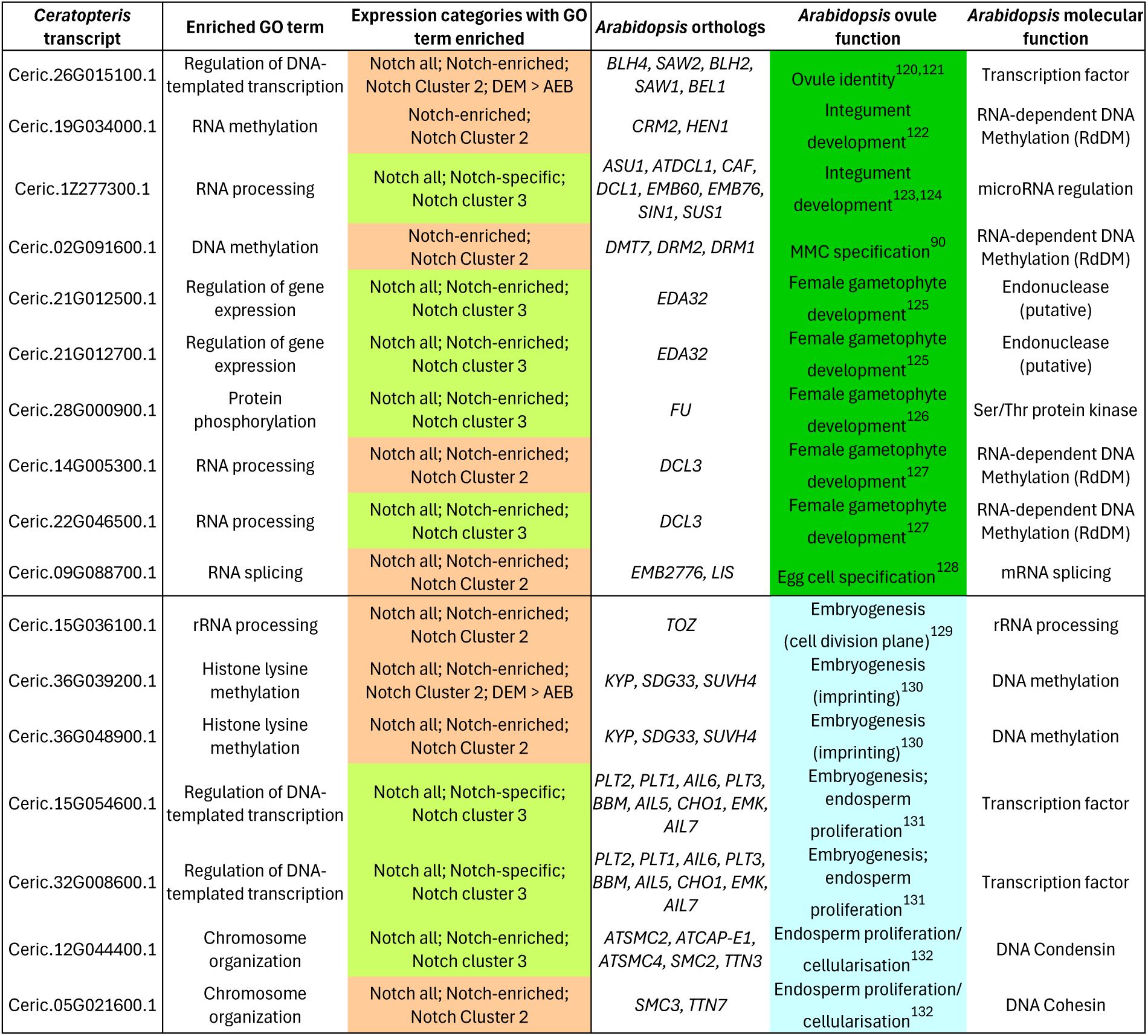
*Ceratopteris* notch reproductive candidate genes enriched in the embryo-bearing archegonium are conserved with *Arabidopsis* ovule/seed genes. List of notch reproductive candidate genes associated with GO terms significantly-enriched within clusters 2 and 3 also orthologous to *Arabidopsis* genes with functions in ovule/seed development. Genes are listed in order of the *Arabidopsis* ortholog’s function during ovule development, separating pre-fertilization events (dark green) from post-fertilization (blue). The two clusters of *Ceratopteris* genes do not correspond to specific stages of ovule/seed development. The enriched GO terms shown represent the most specific term associated with each gene. Full lists of associated GO terms are provided (**Supplementary Data 3**).

**Extended Data Fig. 7.**
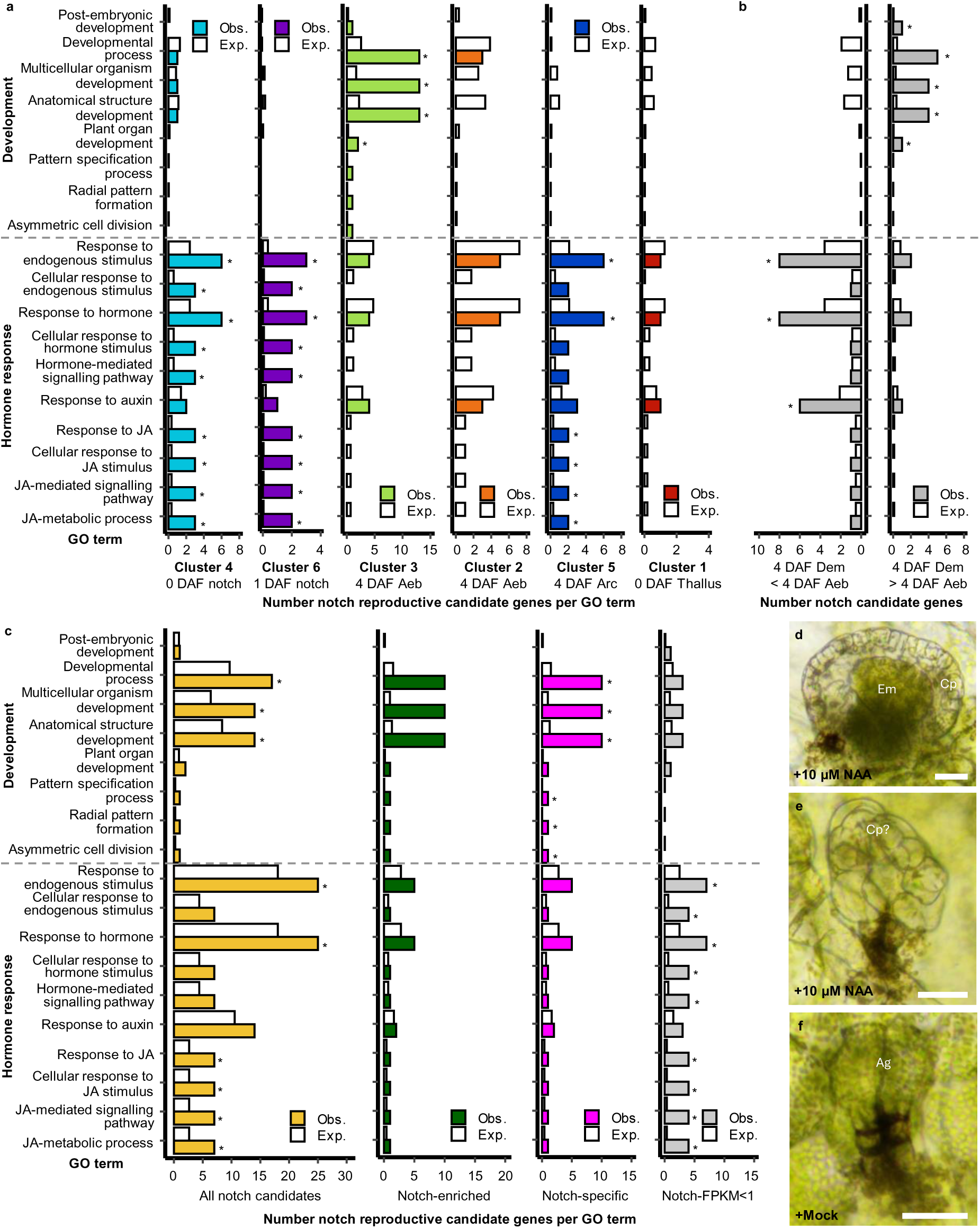
Auxin signalling promotes *Ceratopteris* fertilization-dependent archegonium development. **a-c**, Enrichment of GO terms relating to development or hormone response within each notch expression cluster following the stages of notch development (**a**, see Fig. 5), within genes showing significantly increased or decreased abundance in 4 DAF embryo-bearing archegonia vs 4 DAF dissected embryos (**b**) and within each notch expression category (**c**). Bar plots show the observed (‘Obs.’) and expected (‘Exp.’) gene frequencies per GO term. Asterisks denote a significant increase (p < 0.05) in genes with that GO term compared to that expected based on the whole genome. Individual p-values are provided (**Supplementary Data 3**). **d-f**, Brightfield microscopy of archegonia from embryo-bearing gametophytes after flooding with 10 µM NAA (**d**,**e**) or mock solution (**f**). At 4 DAF embryos were visible inside expanding calyptras under NAA treatment (**d**). Embryo-less archegonia of the same gametophytes exhibited ectopic cell division in response to NAA (**e**) compared to mock-treated controls (**f**). Scale bars = 50 µm. Ag, archegonium, Cp, calyptra, Em, embryo.

**Extended Data Fig. 8.**
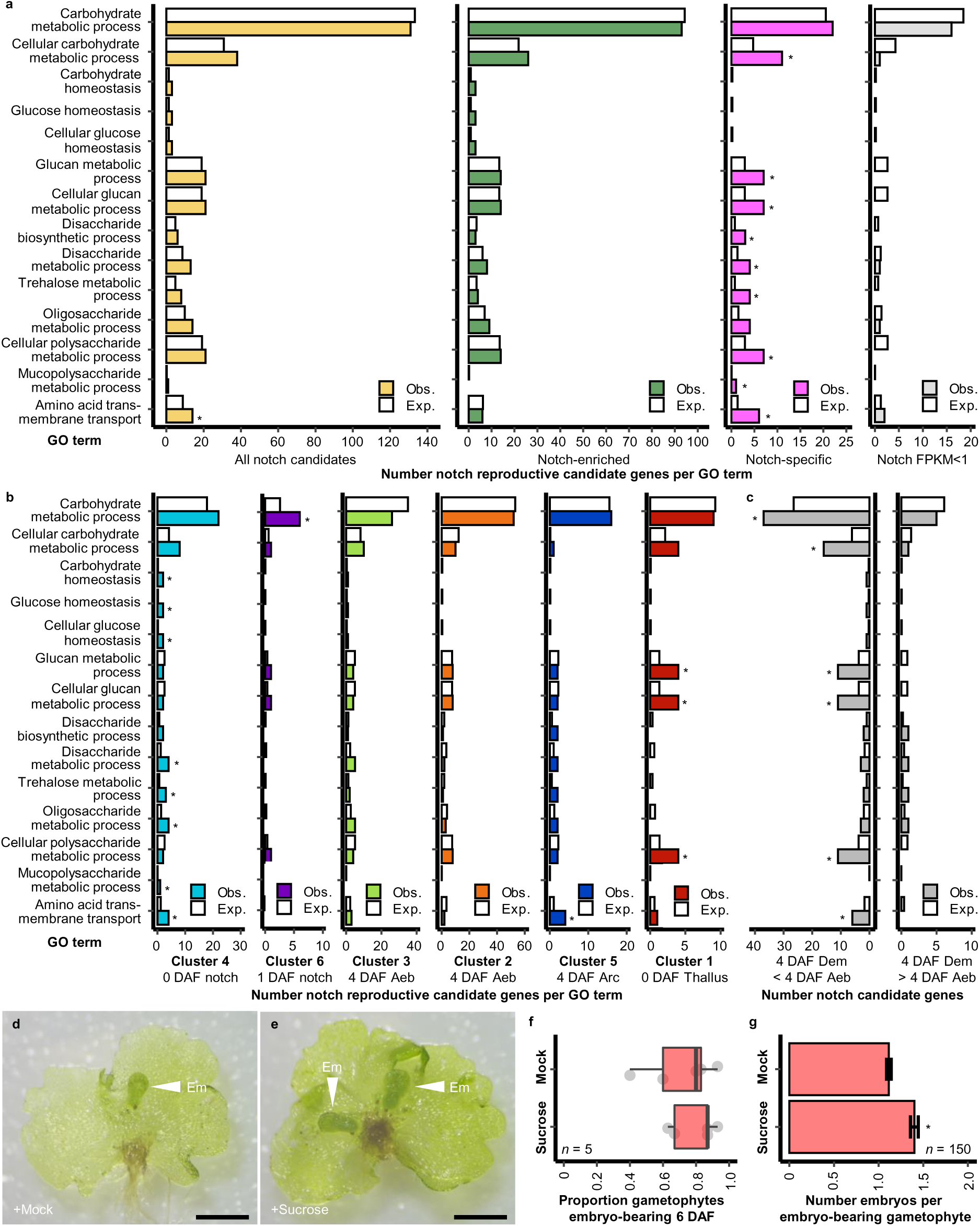
Carbohydrate status plays a role in regulating *Ceratopteris* embryo number. **a-c**, Enrichment of GO terms relating to nutrients (carbohydrate metabolism and amino acid transport) within each notch candidate expression category (**a**), notch expression clusters following the stages of notch development (**b**, see Fig. 5), and within genes showing significantly increased or decreased abundance in 4 DAF embryo-bearing archegonia vs 4 DAF dissected embryos (**c**). Bar plots show the observed (‘Obs.’) and expected (‘Exp.’) gene frequencies per GO term. Asterisks denote a significant increase (p < 0.05) in genes with that GO term compared to that expected based on the whole genome. Individual p-values are provided (**Supplementary Data 3**). **d-g**, Gametophytes typically carry one visible embryo by 6 DAF (**d**), but those flooded with 2% sucrose solution were frequently observed carrying two (**e**). The proportion of gametophytes bearing at least one embryo 6 DAF was similar between mock and sucrose treatments (**f**) whereas the mean number of visible embryos per embryo-bearing gametophyte (**g**) was significantly increased (p < 0.0001). Scale bars = 1 mm. Boxplots denote the median and interquartile range (box) ±1.5x the interquartile range (whiskers). Bar plots denote the mean ± SE. Asterisks denote significant difference (p < 0.05) between mock and sucrose treatment. *n* shown per experiment. Pairwise comparisons were performed using a two-tailed T-test on logit-transformed data (**f**) or a two-tailed Mann-Whitney test (**g**) (**Supplementary Data 4**).

**Extended Data Fig. 9.**
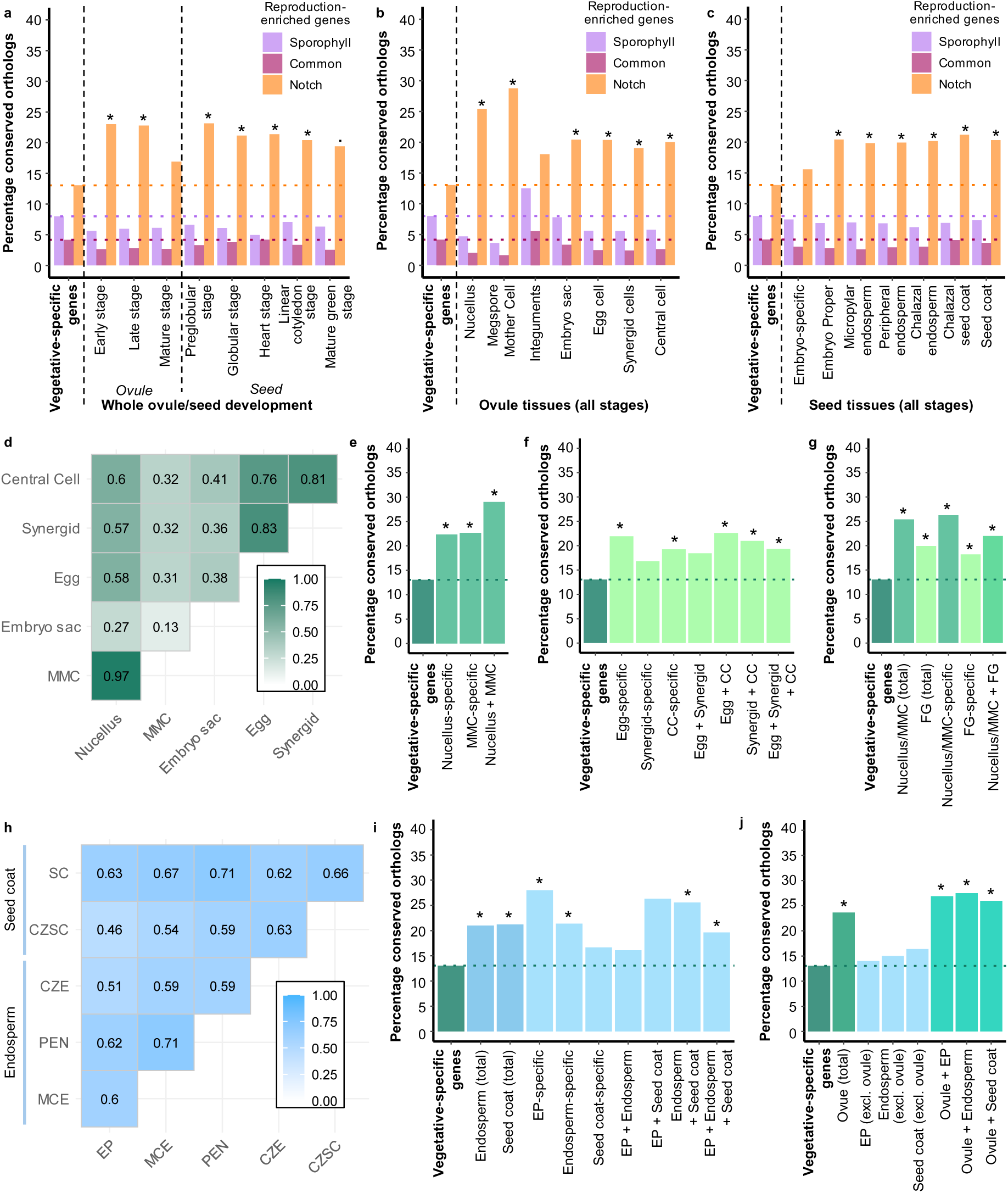
Notch-enriched candidate orthologs overlap with *Arabidopsis* ovule and post-fertilization seed gene networks, but not seed-specific gene networks. **a-c**, Enrichment testing of *Arabidopsis* gene orthologs for sporophyll-enriched, notch-enriched and common orthologs in *Arabidopsis* whole ovule/seed (**a**), ovule tissue (**b**) and seed tissue datasets (**c**) against published vegetative-specific genes. Dotted horizontal lines track the overlap of each candidate ortholog dataset with the vegetative control. **d-j**, Overlap of notch-enriched orthologs significantly-enriched between different tissues within the ovule (**d-g**), within the seed (**h,i**) and between the ovule and seed (**j**). Correlation plots show the Szymkiewicz–Simpson overlap coefficient between each tissue pair (**d,h**). Barplots show enrichment testing of notch-enriched orthologs within genes specific to or common between tissues of the early ovule (**e**), tissues of the female gametophyte (‘FG’) (**f**) the total enriched genes of both (**g**), tissues within the post-fertilization seed (**i**) and all significantly-enriched ovule genes vs seed tissues (**j**). . Asterisks denote a significant increase (Padj < 0.05) in the proportion of overlapping genes between candidate orthologs and the target dataset compared to the overlap with vegetative-specific genes. Individual p-values are provided (**Supplementary Data 8**). The assignment of individual genes to specific tissues is shown in **Supplementary** Fig. 11. EP, embryo proper, MCE, micropylar endosperm, PEN, peripheral endosperm, CZE, chalazal endosperm, CZSC, chalazal seed coat, SC, seed coat

**Extended Data Fig. 10.**
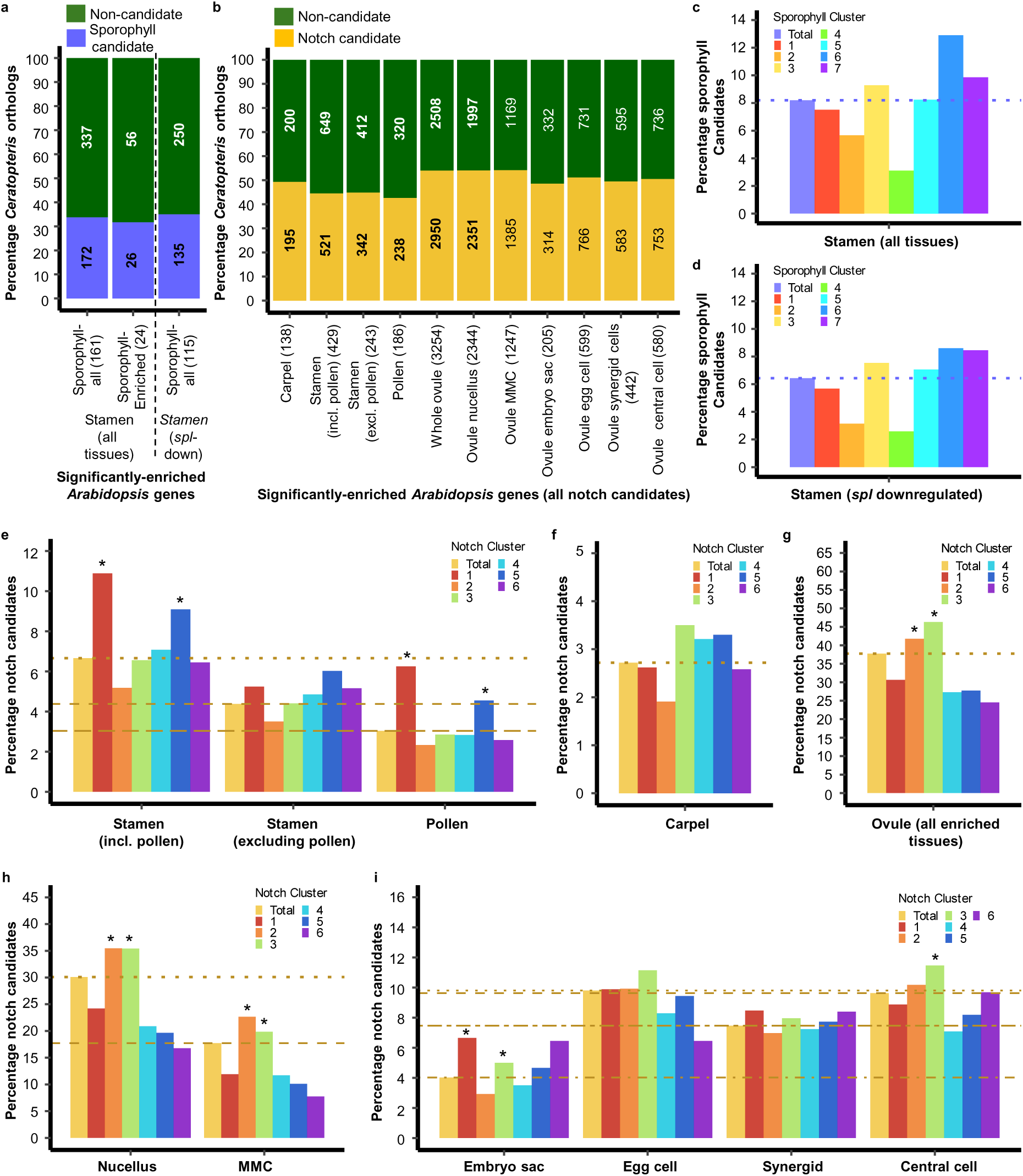
*Ceratopteris* orthologs of conserved *Arabidopsis* ovule genes are significantly enriched in the embryo-bearing archegonium. **a,b**, Reciprocal identification of all *Ceratopteris* orthologs of *Arabidopsis* genes conserved between sporophyll (**a**) or notch candidates (**b**) and significantly-enriched in *Arabidopsis* reproductive organs/tissues (see Fig. 6). The number of conserved *Arabidopsis* genes per dataset is given in brackets. Plots show the proportion of *Ceratopteris* orthologs that are reproductive candidates in the sporophyll (**a**, blue) and notch (**b**, yellow) vs non-candidate genes (green). The number of *Ceratopteris* orthologs within each category is shown. **c,d**, Enrichment testing within all *Ceratopteris* sporophyll candidate genes by expression cluster (see Fig. 3), testing *Ceratotperis* sporophyll candidate orthologs of conserved *Arabidopsis* stamen genes from all stamen tissues (**c**) or those specifically downregulated in *spl* mutant stamens (**d**). **e-i**, Enrichment testing within all *Ceratopteris* notch candidate genes by expression cluster (see Fig. 5), testing *Ceratopteris* notch candidate orthologs of conserved *Arabidopsis* genes in the stamen (**e**), carpel (**f**), whole ovules (**g**) and individual tissues (**h-i**). Asterisks denote a significant increase (Padj < 0.05) in the proportion of genes overlapping between candidate orthologs and each cluster compared to the overlap with all sporophyll or notch candidates, respectively. Individual p-values are provided (**Supplementary Data 9**).

