## Supplementary Figures 1-13 for "Gene networks are conserved across reproductive development between the fern *Ceratopteris richardii* and the flowering plant *Arabidopsis thaliana*"

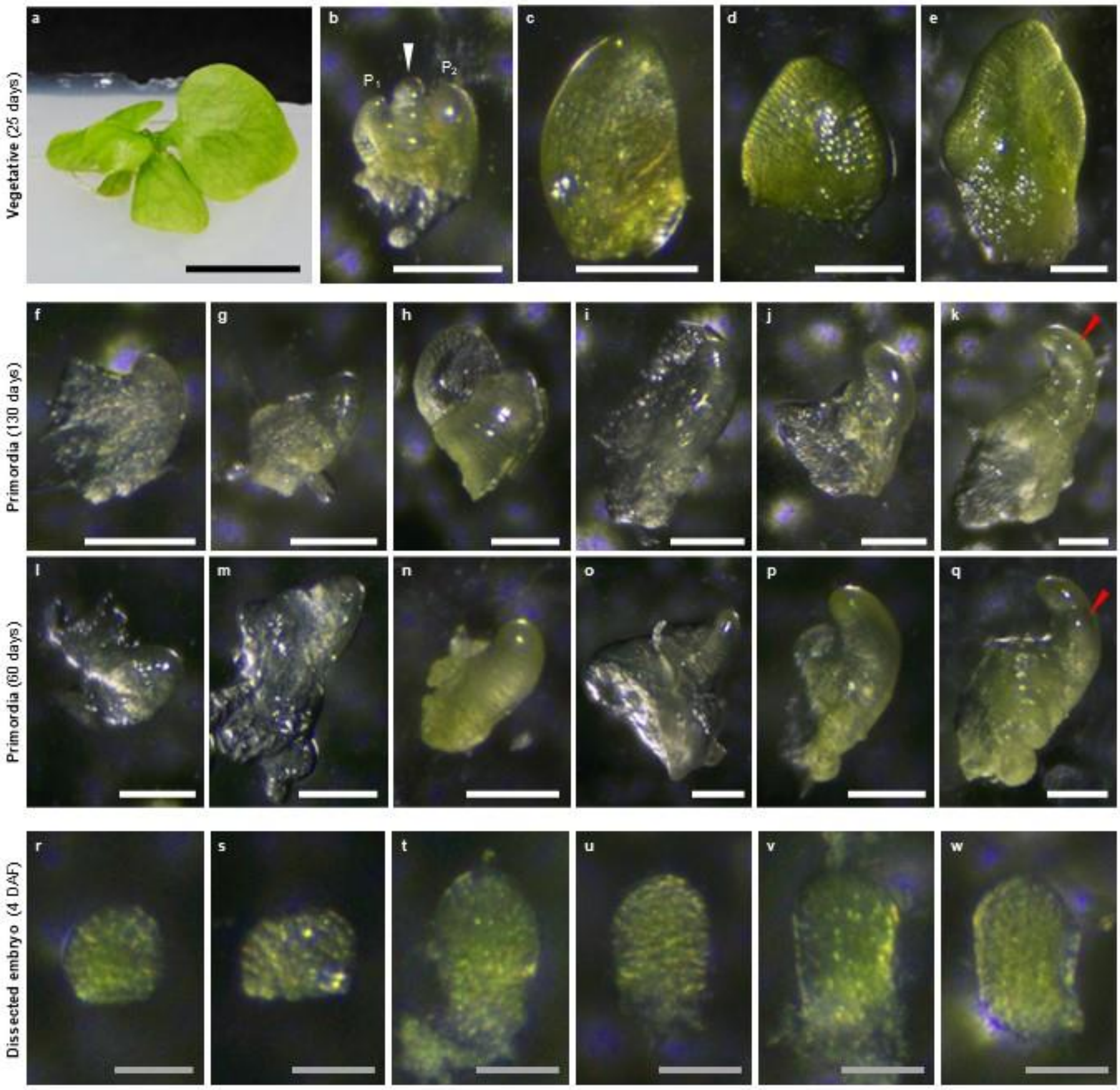


**Supplementary Fig. 1. Additional mRNA-seq samples and developmental ranges.**

**a-e**, At 25 days old *Ceratopteris* sporophytes generate fronds with a simple (spade-like) morphology (**a**). Two samples were dissected for inclusion in mRNA-seq analysis: the shoot apical region (Apx) (**b**) comprising the shoot apex (arrowhead) and recently-emerged primordia ($P_1$, $P_2$), and the earliest dissectible frond primordium within the enclosed shoot apical region (Prm), encompassing a range of sizes (**c-e**).

**f-q**, Primordium developmental stages encompassed by Prm samples at 130d (**f-k**) and 60d (**l-q**). Development within samples at both ages ranged from initial organ outgrowth to curling and the initiation of lateral pinnae (red arrowheads) but excluded complete fiddleheads.

**r-w**, Embryo development stages encompassed by dissected 4 DAF dissected embryo (Dem) samples.

Scale bars = 10mm (black), 200 μm (white) and 100 μm (grey).

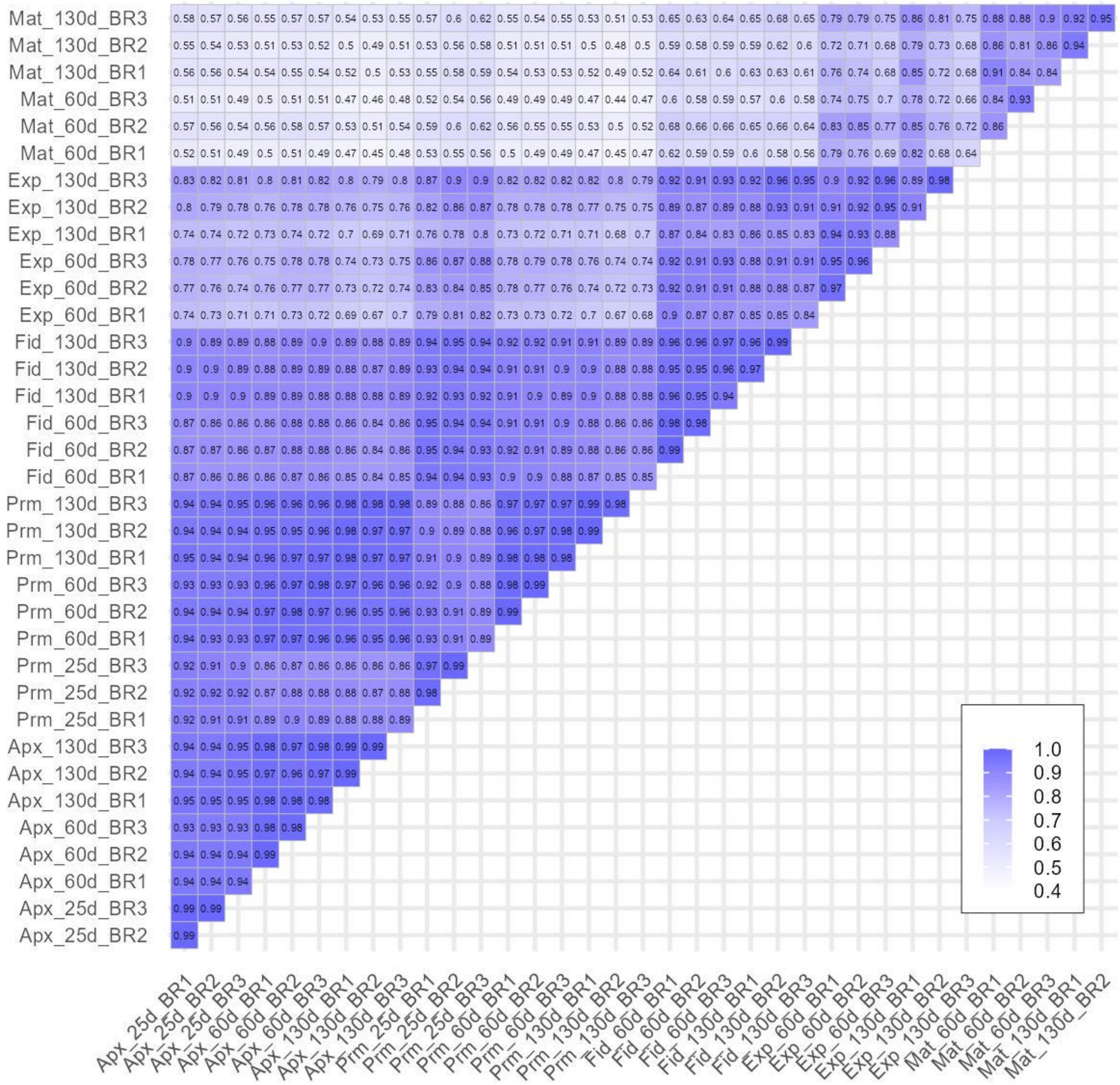


**Supplementary Fig. 2. Correlation analysis between all sporophyte mRNA-seq samples.** Samples are grouped by biological replicate, and then within each frond developmental stage sampled in order of frond development. Colour intensity denotes increasing correlation, as shown. Samples typically show the highest correlation between biological replicates, and then between vegetative and reproductive samples within the same frond developmental stage, with the exception of 25d primordia which correlate more strongly with the fiddlehead stage than other primordia. Correlations between developmental stages typically decline progressively with developmental distance.

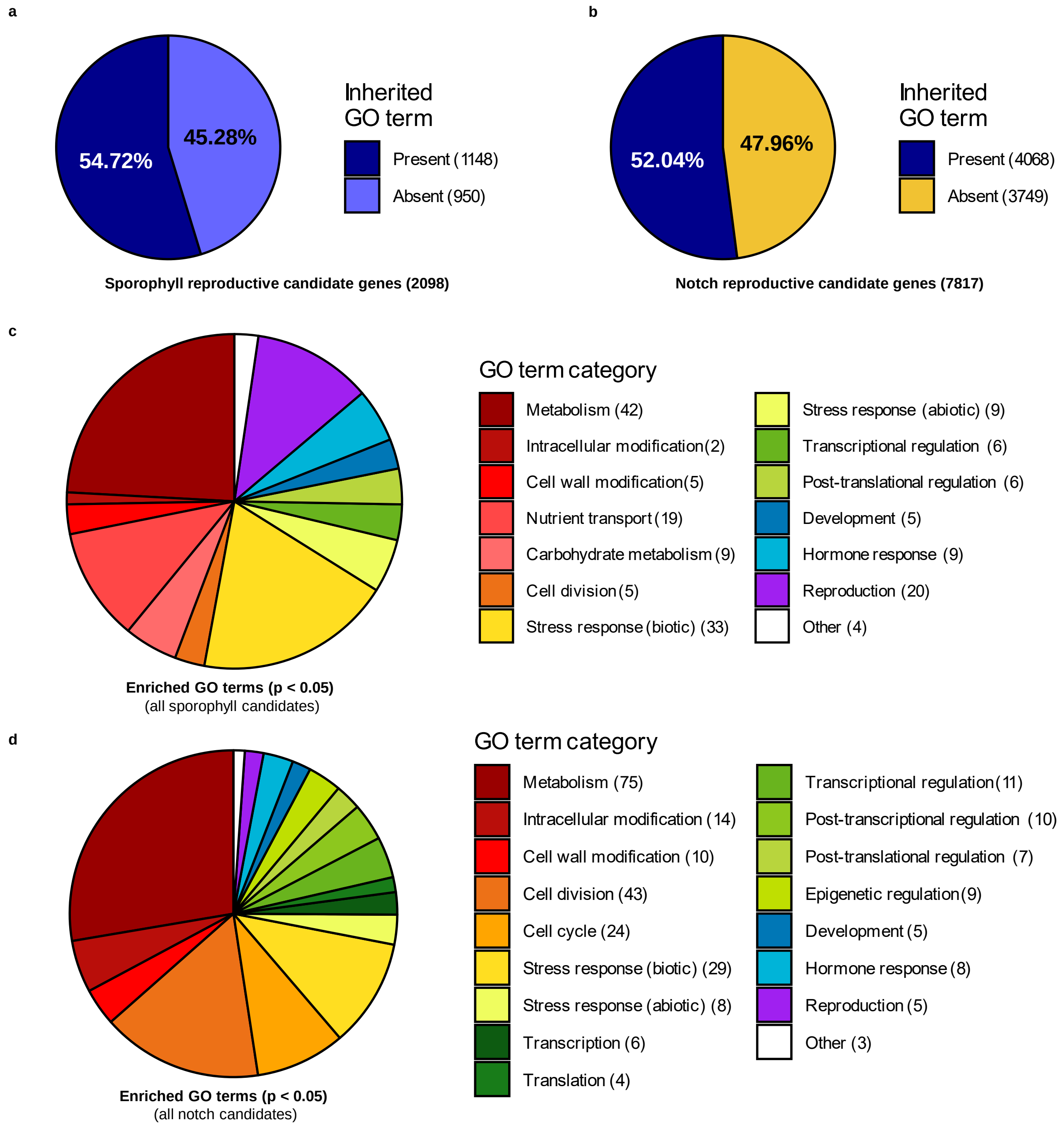


**Supplementary Fig. 3. Functional categorisation of enriched GO terms in sporophyll and notch reproductive candidate genes.**

**a**,**b**, The proportion of *Ceratopteris* genes with GO terms inherited from orthologous genes within all sporophyll reproductive candidates (**a**) and all notch candidates (**b**).

**c**,**d** The proportion of different GO term categories significantly enriched ($p < 0.05$) within all sporophyll reproductive candidates (**c**) and all notch reproductive candidates (**d**). Numbers in brackets within the key correspond to the numbers of GO terms within each category.

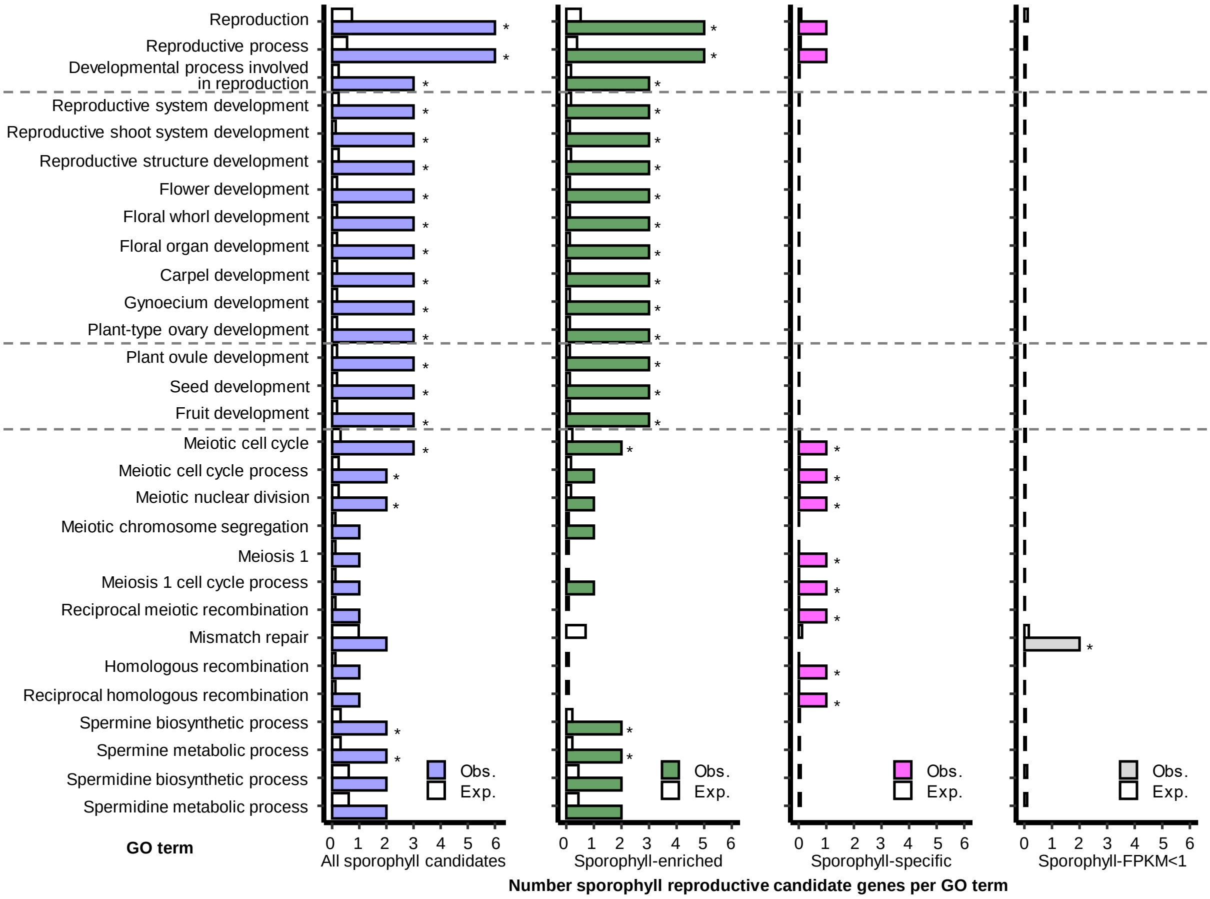


**Supplementary Fig. 4. Distribution of enriched reproductive GO terms across sporophyll reproductive candidate expression categories.**

Frequency of observed sporophyll reproductive candidate genes with individual reproductive GO terms against the frequency expected based on the whole genome, comparing between all sporophyll candidates, sporophyll-enriched candidates, sporophyll-specific candidates and sporophyll-FPKM<1 candidates, as shown.

All reproductive GO terms in which significant enrichment was detected within the sporophyll are shown. Obs., observed frequency, Exp., expected frequency. Asterisks denote a significant increase ($p < 0.05$) in the frequency of genes with that GO term compared to the expected frequency.

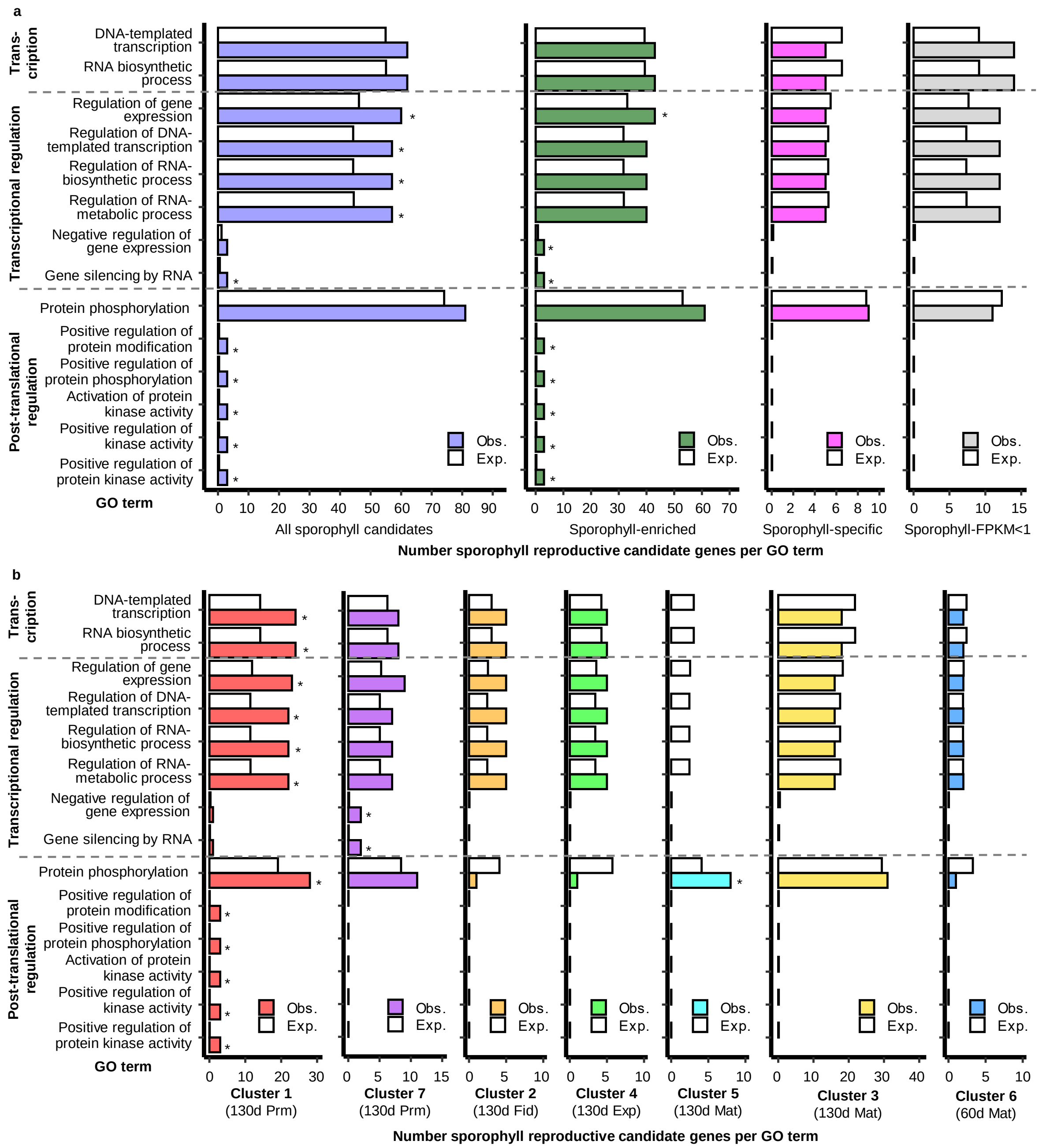


**Supplementary Fig. 5. Distribution of enriched gene regulatory GO terms across sporophyll reproductive candidate expression categories/clusters.**

The frequency of observed sporophyll reproductive candidate genes with individual GO terms relating to development or hormone regulation against the frequency expected based on the whole genome, analysed within sporophyll expression categories (**a**) and within each expression cluster (**b**) in order of their peak in abundance during sporophyll development (as shown, see **Fig. 3b**).

All reproductive GO terms in which significant enrichment was detected within the sporophyll are shown. Obs., observed frequency, Exp., expected frequency. Asterisks denote a significant increase ($P < 0.05$) in the frequency of genes with that GO term compared to the expected frequency.

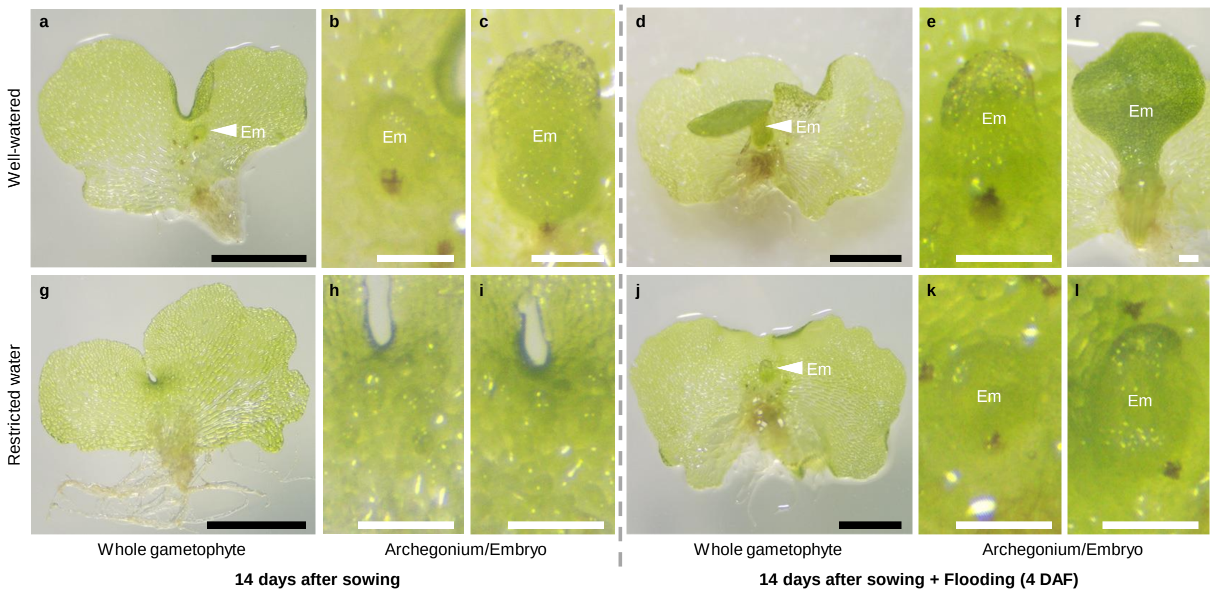


**Supplementary Fig. 6. *Ceratopteris* embryo development is synchronised through restricted watering.**

Comparison of gametophyte and embryo morphology 14 days after sowing between well-watered (**a-f**) and restricted water growth conditions (**g-l**) in unflooded populations 14 days after sowing (**a-c**, **g-i**) and 4 days subsequently after flooding (**d-f**, **j-l**). In well-watered populations, embryos (Em) were already visible on gametophytes at 14 days after sowing (**a**) with a range of different developmental stages detectable (**b,c**). This disparity persisted in the population after flooding (**d-f**). Under restricted water, embryos were not visible 14 days after sowing (**g-i**) and after flooding the developmental range of visible embryos was much closer (**j-l**). Scale bars = 1 mm (black), 200 µm (white).

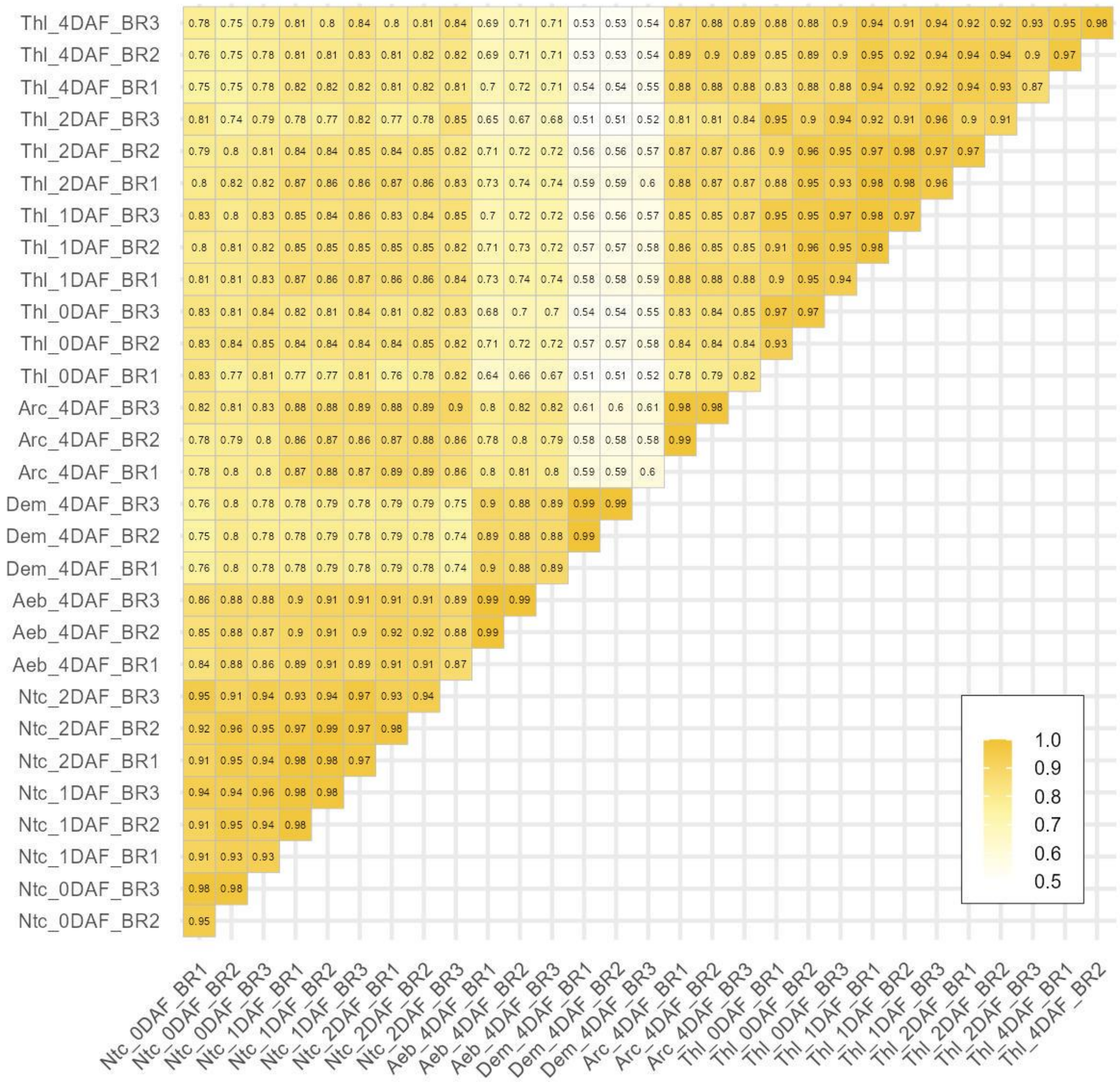

**Supplementary Fig. 7. Correlation analysis between all gametophyte mRNA-seq samples.**

Samples are grouped by biological replicate, and then within each developmental stage within notch (Ntc) and thallus (Thl) groups. Colour intensity denotes increasing correlation, as shown. Samples typically show the highest correlation between biological replicates, and then within notch and thallus groups of samples, with the exception dissected embryos (Dem), which correlate more closely with embryo-bearing archegonia (Aeb) than other samples.

**a**

| Protein | Species | E-value | % identity | Align len | Strands | Query ID | Query from | Query to | Target fron | Target to | Bitscore | # identical | Positives | Gaps | Query len | Target len |
|---|---|---|---|---|---|---|---|---|---|---|---|---|---|---|---|---|
| Ceric.31G036600.1.p | C.richardii v2.1 | 0 | 99 | 840 | +/+ | CrANT_Bui_et_al_2017 | 1 | 838 | 113 | 951 | 1671.75 | 829 | 829 | 3 | 838 | 952 |
| Ceric.31G036600.4.p | C.richardii v2.1 | 0 | 99 | 811 | +/+ | CrANT_Bui_et_al_2017 | 30 | 838 | 7 | 816 | 1612.43 | 799 | 800 | 3 | 838 | 817 |
| Ceric.31G036600.3.p | C.richardii v2.1 | 0 | 99 | 797 | +/+ | CrANT_Bui_et_al_2017 | 44 | 838 | 1 | 796 | 1580.46 | 786 | 786 | 3 | 838 | 797 |
| Ceric.31G036600.2.p | C.richardii v2.1 | 0 | 94 | 840 | +/+ | CrANT_Bui_et_al_2017 | 1 | 838 | 113 | 912 | 1568.13 | 790 | 790 | 42 | 838 | 913 |
| Ceric.12G002600.1.p | C.richardii v2.1 | 1.07E-124 | 49 | 550 | +/+ | CrANT_Bui_et_al_2017 | 355 | 837 | 516 | 1026 | 401.364 | 268 | 315 | 106 | 838 | 1028 |
| Ceric.10G056200.1.p | C.richardii v2.1 | 2.07E-105 | 81 | 218 | +/+ | CrANT_Bui_et_al_2017 | 357 | 571 | 444 | 660 | 347.051 | 176 | 189 | 4 | 838 | 915 |
| Ceric.10G056200.2.p | C.richardii v2.1 | 2.68E-105 | 81 | 218 | +/+ | CrANT_Bui_et_al_2017 | 357 | 571 | 422 | 638 | 346.28 | 176 | 189 | 4 | 838 | 893 |
| Ceric.26G003300.1.p | C.richardii v2.1 | 2.97E-102 | 80 | 215 | +/+ | CrANT_Bui_et_al_2017 | 357 | 568 | 303 | 513 | 332.798 | 172 | 185 | 7 | 838 | 683 |
| Ceric.32G008600.3.p | C.richardii v2.1 | 1.54E-101 | 58 | 320 | +/+ | CrANT_Bui_et_al_2017 | 257 | 568 | 67 | 373 | 330.102 | 187 | 225 | 21 | 838 | 665 |
| Ceric.32G008600.2.p | C.richardii v2.1 | 2.05E-99 | 58 | 320 | +/+ | CrANT_Bui_et_al_2017 | 257 | 568 | 339 | 645 | 331.643 | 187 | 225 | 21 | 838 | 937 |
| Ceric.32G008600.1.p | C.richardii v2.1 | 2.71E-99 | 58 | 320 | +/+ | CrANT_Bui_et_al_2017 | 257 | 568 | 345 | 651 | 331.257 | 187 | 225 | 21 | 838 | 943 |

**b**

| Protein | Species | E-value | % identity | Align len | Strands | Query ID | Query from | Query to | Target fron | Target to | Bitscore | # identical | Positives | Gaps | Query len | Target len |
|---|---|---|---|---|---|---|---|---|---|---|---|---|---|---|---|---|
| Ceric.33G031700.1.p | C.richardii v2.1 | 0 | 100 | 370 | +/+ | CrLFY1_Plackett_et_al_2018 | 1 | 370 | 112 | 481 | 747.658 | 370 | 370 | 0 | 370 | 482 |
| Ceric.33G031700.2.p | C.richardii v2.1 | 0 | 100 | 370 | +/+ | CrLFY1_Plackett_et_al_2018 | 1 | 370 | 100 | 469 | 747.273 | 370 | 370 | 0 | 370 | 470 |
| Ceric.33G031700.3.p | C.richardii v2.1 | 0 | 100 | 370 | +/+ | CrLFY1_Plackett_et_al_2018 | 1 | 370 | 1 | 370 | 745.732 | 370 | 370 | 0 | 370 | 371 |
| Ceric.18G076300.3.p | C.richardii v2.1 | 0 | 85 | 379 | +/+ | CrLFY1_Plackett_et_al_2018 | 1 | 370 | 17 | 394 | 650.588 | 323 | 342 | 10 | 370 | 395 |
| Ceric.18G076300.1.p | C.richardii v2.1 | 0 | 85 | 379 | +/+ | CrLFY1_Plackett_et_al_2018 | 1 | 370 | 17 | 394 | 650.588 | 323 | 342 | 10 | 370 | 395 |
| Ceric.33G031700.4.p | C.richardii v2.1 | 3.22E-179 | 98 | 251 | +/+ | CrLFY1_Plackett_et_al_2018 | 120 | 370 | 11 | 261 | 498.819 | 246 | 249 | 0 | 370 | 262 |
| Ceric.18G076300.2.p | C.richardii v2.1 | 8.84E-164 | 90 | 247 | +/+ | CrLFY1_Plackett_et_al_2018 | 124 | 370 | 5 | 251 | 459.144 | 222 | 229 | 0 | 370 | 252 |
| Ceric.18G076300.4.p | C.richardii v2.1 | 3.48E-157 | 91 | 235 | +/+ | CrLFY1_Plackett_et_al_2018 | 136 | 370 | 1 | 235 | 441.81 | 213 | 220 | 0 | 370 | 236 |

**Supplementary Fig. 8. Validation of notch reproductive candidate genes as previously-published embryo-related genes.**

**a,b**, BLASTp search results of *C. richardii* genome v2.1 against the amino acid sequences of CrANT (**a**) and CrLFY1 (**b**), showing the genomic loci with the closest identity. Closest matches were identified as Ceric.31G036600 and Ceric.33G031700, respectively.

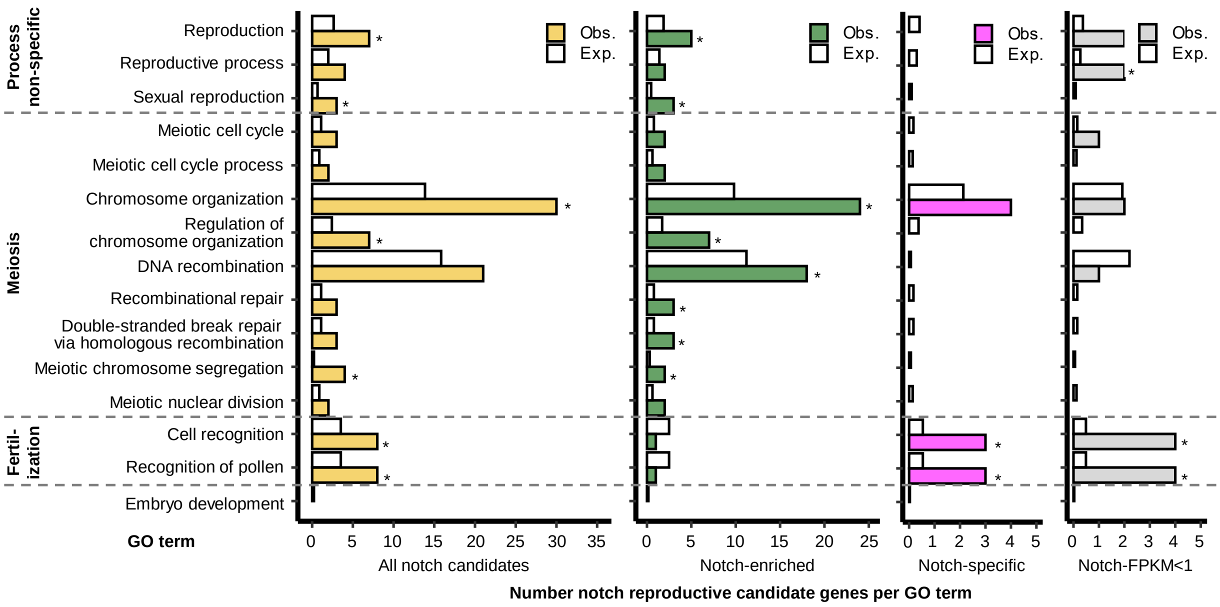


**Supplementary Fig. 9. Distribution of enriched reproductive GO terms across notch reproductive candidate expression categories.**

Frequency of observed notch reproductive candidate genes with individual reproductive GO terms against the frequency expected based on the whole genome, comparing between all notch candidates, notch-enriched candidates, notch-specific candidates and notch-FPKM<1 candidates, as shown.

All reproductive GO terms in which significant enrichment was detected within notch reproductive candidates are shown. Obs., observed frequency, Exp., expected frequency. Asterisks denote a significant increase ($p < 0.05$) in the frequency of genes with that GO term compared to the expected frequency.

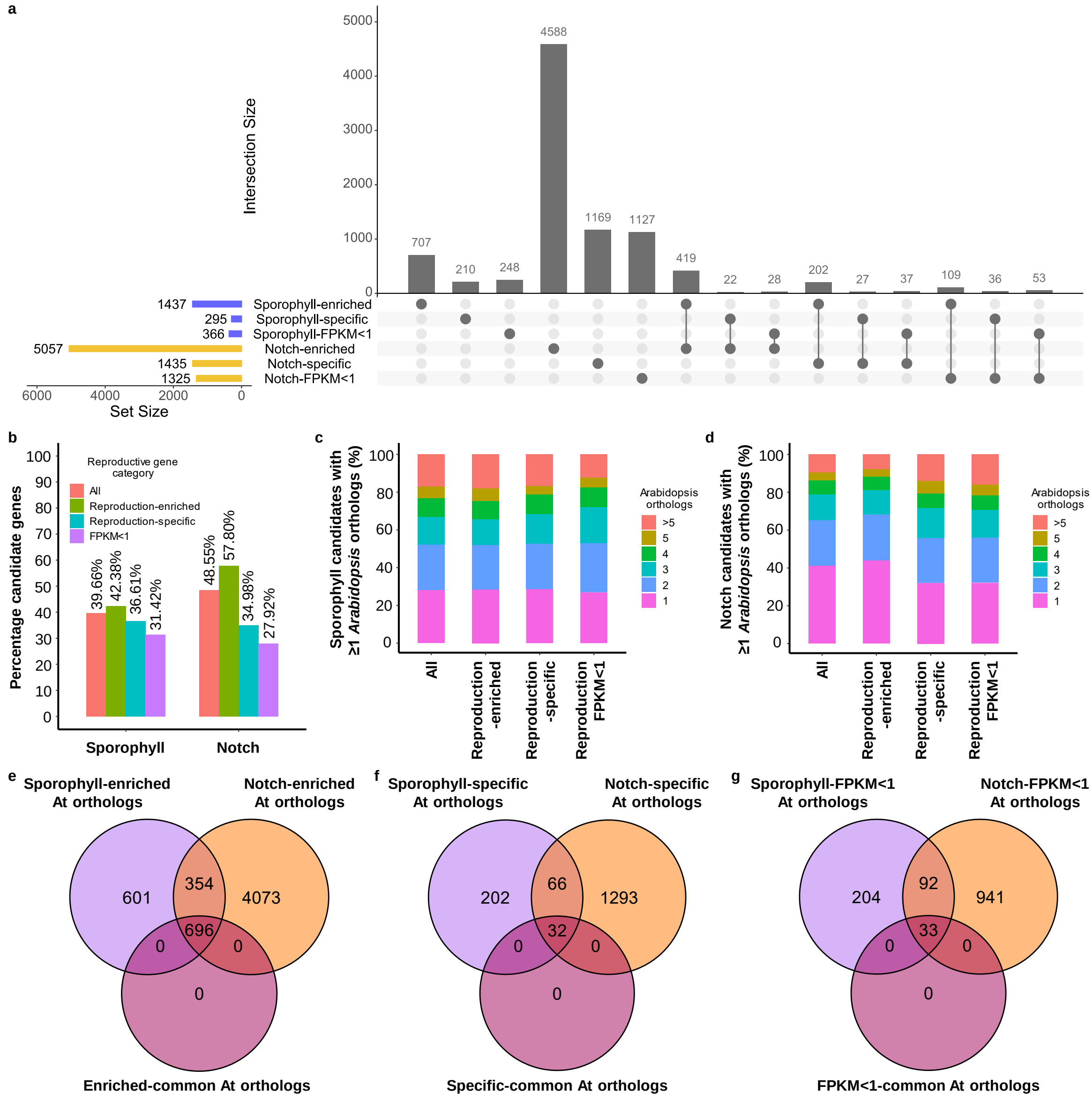


**Supplementary Fig. 10. *Arabidopsis* gene orthology to *Ceratopteris* reproductive candidates by expression category.**

**a**, Upset plot of reproductive candidate genes in the *Ceratopteris* sporophyll (blue) and notch (yellow) by expression category (see **Fig. 1n, 4j**), showing the frequency of overlap between each organ + category combination.

**b-d**, *Arabidopsis* orthology to *Ceratopteris* candidate genes by expression category, showing the percentage of candidate genes with direct *Arabidopsis* orthologs (**b**) and, within those genes, the proportion of candidates with differing specificity of *Arabidopsis* orthologs within the sporophyll (**c**) and the notch (**d**).

**e-g**, Frequency and overlap of nonredundant *Arabidopsis* orthologs of sporophyll, notch and common candidates within reproduction-enriched (**e**), reproduction-specific (**f**) and FPKM<1 expression categories (**g**).

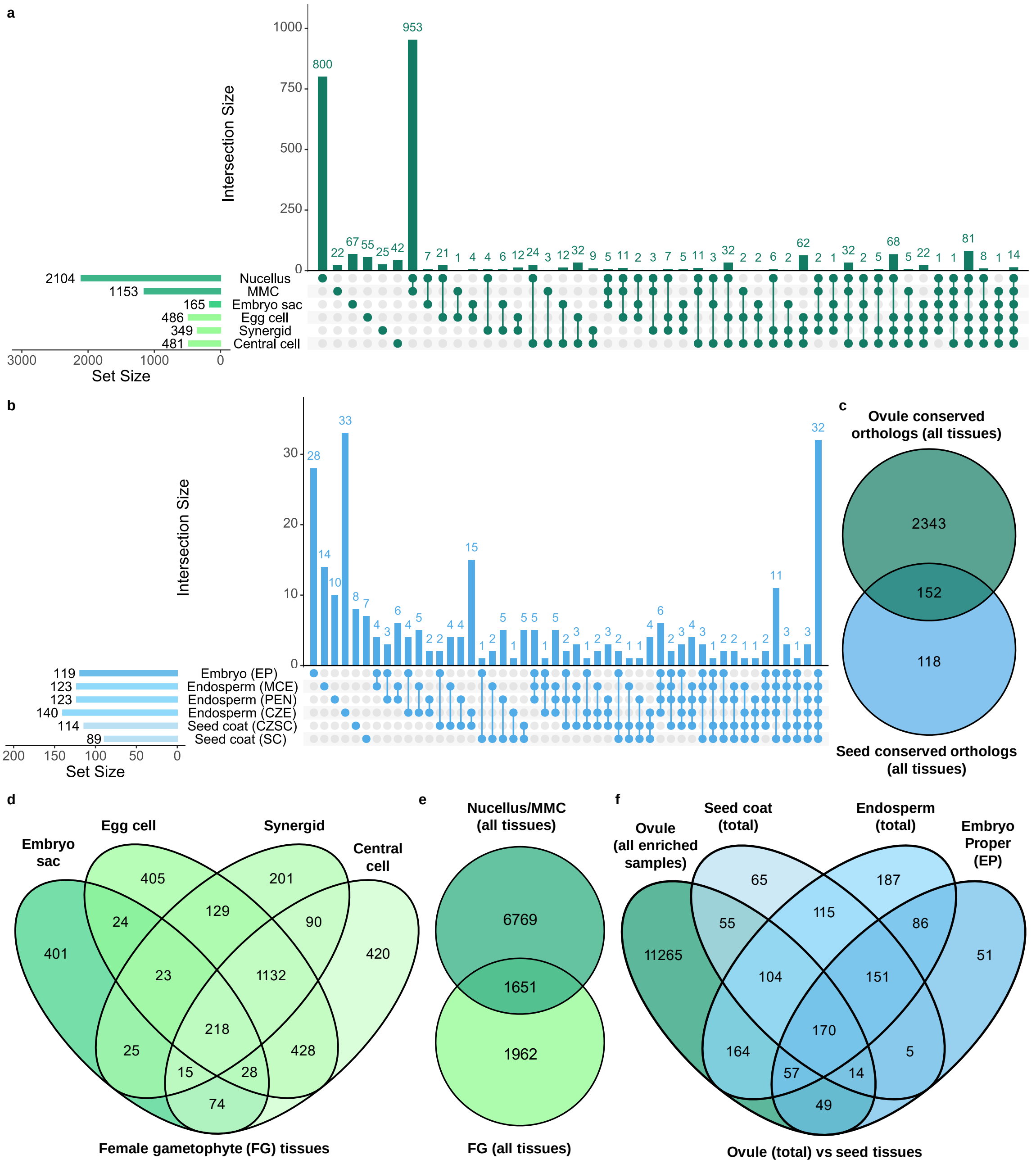


**Supplementary Fig. 11. Comparison of significantly enriched notch-enriched *Arabidopsis* orthologs between ovule and seed tissues.**

**a,b,** Upset plot of significantly-enriched *Arabidopsis* orthologs of notch-enriched candidates (see **Extended Data Fig. 9**) between tissues within the ovule (**a**) and seed (**b**). Different shades in (**a**) denote datasets obtained from separate studies (see **Fig. 6**) and in (**b**) denote separate tissues of origin (embryo, endosperm or seed coat).

**c**, Venn comparison of the *Arabidopsis* orthologs of notch-enriched candidates significantly enriched ($P_{adj} < 0.05$) in ovule and seed tissues, totalling all significantly-enriched datasets.

**d-f**, Venn comparisons of all *Arabidopsis* reproductive genes published between tissues of the female gametophyte (**d**), between tissue groups within the ovule (**e**) and between seed tissues (totalled by tissue of origin) vs all reproductive genes in the ovule (**f**).

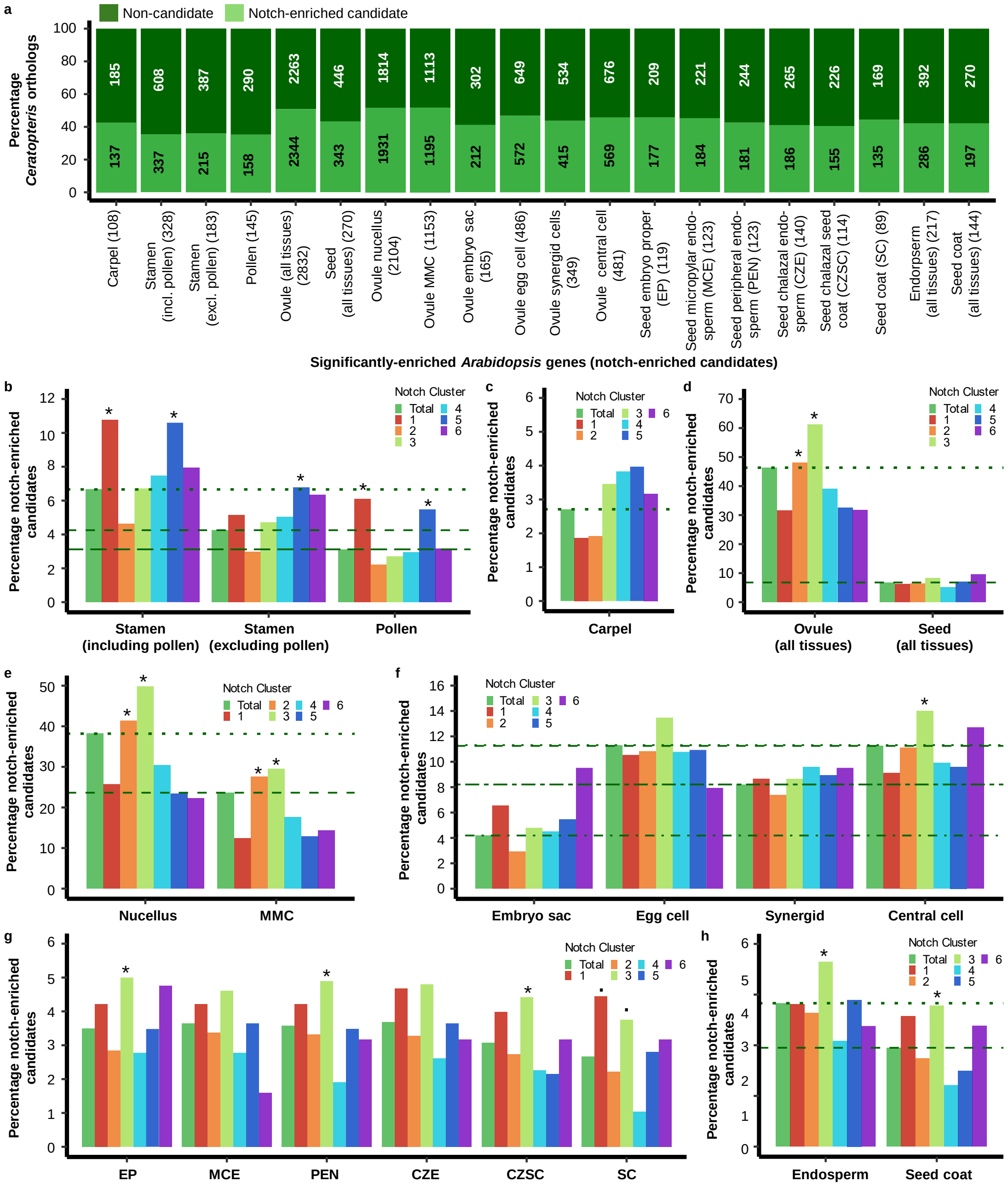


**Supplementary Fig. 12. Reciprocal enrichment testing of *Ceratopteris* notch-enriched orthologs of conserved *Arabidopsis* genes within notch-enriched candidates.**

**a**, Reciprocal identification of all *Ceratopteris* notch-enriched orthologs of conserved *Arabidopsis* reproductive genes by *Arabidopsis* organ/tissue, as in **Extended Data Fig. 9**.

**b-h**, Enrichment testing of conserved candidate genes by notch expression cluster (as shown) within notch-enriched candidates only. 'Endosperm' and 'Seed coat' were calculated as a nonredundant total of conserved genes in individual endosperm tissues (MCE, PEN, CZE) and seed coat tissues (CZSE, SC), respectively. Asterisks denote a significant increase ($P_{adj}$ < 0.05) in the proportion of candidate orthologs overlapping in a cluster compared with all candidates ('total'). '·' Denotes marginal significance ($P_{adj}$ = 0.05). Individual p-values provided (**Supplementary Data 9**).

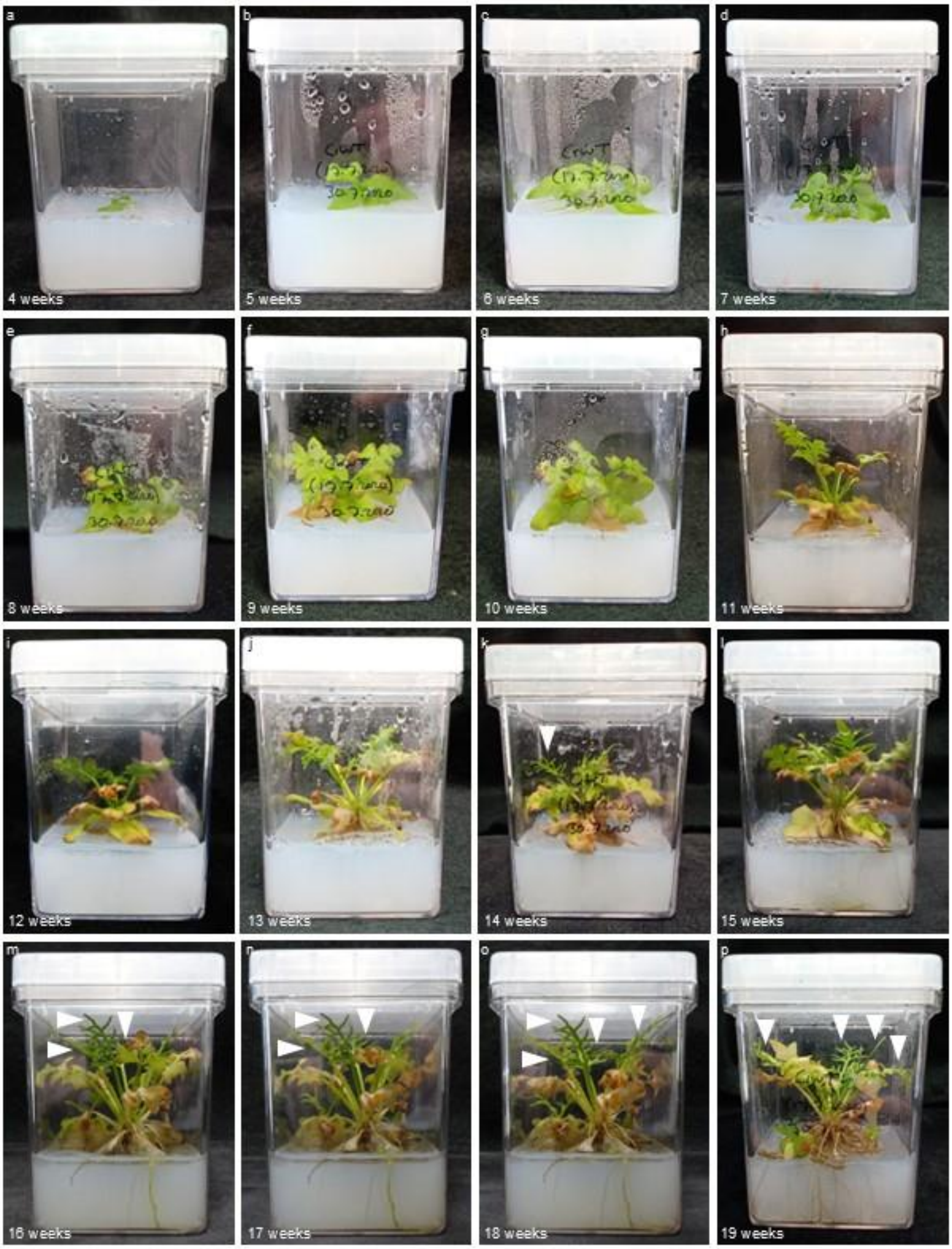


**Supplementary Figure 13. Growth of the *Ceratopteris* sporophyte under tissue culture conditions.**

**a-p**, Developmental series of the *Ceratopteris* sporophyte grown under sterile tissue-culture for reproductive characterisation, shown at weekly intervals from four weeks old (**a**) until tissue harvesting for mRNA-seq analysis at 19 weeks (**p**). Sporophylls are morphologically distinct and discernible from preceding vegetative fronds by eye (white arrowheads).
